# EPHA2-Ephrin-B1 *cis*-interaction supports self-renewal ability leading to recurrence of oral cancer

**DOI:** 10.1101/2025.01.03.631197

**Authors:** Reshma Raj R, Nandini Datta, Soumya Krishnan U, Geetha Shanmugam, Jiss Maria Louis, Ravichandran Damodaran Jeyaram, Keshava K Datta, Harsha Gowda, Meghna Sarkar, Madhumathy G Nair, Jyothy S Prabhu, Hafsa Shabeer, Balagopal P G, Thameem A, Alan Jose, Vishnu Sunil Jaikumar, Tessy Thomas Maliekal

## Abstract

Eph-Ephrin pathway that drives a bidirectional signaling regulates a plethora of biological activities with its varied level of complexity. While the canonical *trans*-interaction of the ligand and receptor on neighboring cells initiates forward signaling that brings about biological activities, their *cis*-interaction on the same cell attenuates the forward signaling, modulating the biological effects. Yet, non-canonical *cis*-interactions with heterotypic surface proteins and kinases regulate certain biological effects. In cancer, the canonical signaling is believed to be tumor suppressive, while the ligand-independent non-canonical signaling drives tumor progression and poor prognosis. Self-renewal ability of cancer cells is a major underlying cause of recurrence and poor prognosis of cancer. Using SILAC-based proteomics, we identified Ephrin-B1 signaling as a crucial regulator of oral cancer self-renewal. Though Ephrin-B1 is known to regulate normal stem cells, its role in cancer remains underexplored. Our biochemical analyses show that Ephrin-B1 binds to a nonconventional receptor EPHA2, which is known to regulate cancer stem cells (CSCs), though the mechanism is less explored. Contrary to the belief that the *cis*-interaction of receptors and ligands is a means to block functional signaling, our immunoprecipitation, FRET facilitated photoswitching analysis, proximity ligation assay, and *in vitro* kinase assay provide evidence for the Ephrin-B1-EPHA2 *cis*-interaction leading to the phosphorylation of EphrinB1 at Y324/329 and Y317. Extreme limiting dilution assay *in vitro* and *in vivo* confirmed that this *cis*-interaction promotes CSC enrichment. Substantiating our *in vitro* results, mouse orthotopic models showed that Ephrin-B1/EPHA2 interaction regulates prognosis. The clinical relevance of the finding was validated using a TCGA data set and immunohistochemical analysis of tissue microarray using samples from oral cancer patients with recurrence in comparison to patients, who showed disease-free survival. Given that Ephrin-B interacts with EphB for normal stem cell homeostasis, this unconventional EPHA2/Ephrin-B1 *cis*-interaction, specifically manifested in CSC niches, might serve as an attractive target for therapy, warranting further validation.

## Introduction

Eph/Ephrin signaling is a bidirectional signaling, which is mediated through membrane-bound Eph receptors and Ephrin ligands. Eph receptors are tyrosine kinases, which are classified into A or B based on the similarity in the sequence. Humans have nine EphA receptors, five EphB receptors, five Ephrin-A ligands with glycosylphosphatidylinositol (GPI) linkage, and three Ephrin-B ligands with a transmembrane region. Eph/Ephrin signaling contributes to varied context-dependent functions *via* a diverse range of receptor-ligand combinations, bidirectional or unidirectional signaling, and Ephrin-independent non-canonical signaling of Eph receptors or Eph-independent signaling of Ephrin ligands (1). During development, it is known to regulate axon growth (2), tissue patterning (3), tissue boundary formation (4), migration of stem cells (5, 6), and stem cell maintenance (7–10). Complementary to these developmental processes, several tumorigenic properties like, proliferation, apoptosis, angiogenesis, metastasis, and immune modulation, are also regulated by the Eph/Ephrin pathway (1). Thus, it is evident that unraveling the complexity of this evolutionarily conserved pathway is critical for understanding different biological processes.

In a typical bidirectional signaling, when the receptor molecule of one cell interacts with the ligand molecule on a neighboring cell, i.e., a *trans* interaction, it leads to signaling events in the receptor-bearing cell (forward signaling) and the ligand-bearing cell (reverse signaling). In this scenario, we would expect the ligand and the receptor to be present on the neighboring cells. But in many contexts, it is shown that cells co-express the ligands and the receptors, allowing the possibility of a *cis*-interaction, i.e., interaction within the same cell. Using an overexpression system, Yin *et al* have shown that the *cis*-interaction of EphA/Ephrin-A does not lead to signaling, but it competes with the *trans* signaling, reducing the tyrosine phosphorylation of the receptor (11). When the role of *cis* or *trans*-interactions of co-expressed Eph receptors and Ephrin ligands in axonal guidance was tested, it was observed that the majority of the ligands and receptors of the same cell segregate to different growth cones, minimizing the chance of *cis*-interaction (12). Eventually, several other studies showed that the balance between *cis* and *trans* modes of ligand-receptor interaction is key to the diversity of axon guidance signaling responses (13). The same notion that *cis*-interaction is a means of regulating the *trans*-interaction, is shown in cancer cells also. One of the reports from Pasquale’s group has shown that the *cis*-interaction of EphA3 with Ephrin-A3 or Ephrin-B2 is a means of regulation of EphA3 forward signaling in lung cancer. Further, they have shown that one of the reported EphA3 mutations in lung cancer patients enhances its *cis*-interaction to suppress the tumor suppressive effects of EphA3 signaling (14). Apart from the *cis* interaction between its own family members, these ligands and receptors can have heterotypic *cis*-interactions with other signaling molecules like epidermal growth factor receptor (EGFR), fibroblast growth factor receptor (FGFR), Dishevelled2 (Dvl2), etc., to modulate several physiological functions, which is extensively reviewed elsewhere (15). Taken together, the receptors of this family can have canonical ligand-dependent activity as well as non-canonical ligand-independent activity. Thus, it is evident that the outcome of Eph-Ephrin signaling depends on the type of ligands and receptors, mode of interaction, and the type of heterotypic interactions. Given the complexity of the system, the downstream pathway in each of these contexts is not resolved. However, phosphorylations of some residues of the receptors and ligands are reported. Yet, there are considerable lacunae in our understanding of how the phosphorylations of the receptors and ligands lead to the specific effect. With its different layers of intricacy, this signaling can tightly regulate developmental processes as well as cancer.

Several members of the Eph-Ephrin family are shown to be involved in regulating various aspects of malignancy. As the field evolved, there is contradicting evidence suggesting the tumor suppressive as well as tumor promoting role of this signaling (16). In general, it is accepted that the canonical forward signaling exerts tumor suppression, while the ligand-independent-non-canonical signaling promotes tumor progression (1). Although deregulations of several of the family members are reported in a wide range of cancers, some of the molecules that emerge as therapeutic targets are EPHA2 and EPHB2 (17–21). These receptors regulate cell proliferation, invasion, metastasis, immune modulation, and cancer stem cell (CSC) properties. EPHB2 has been shown to exert immunomodulation in the tumor microenvironment, and it regulates angiogenesis (16). On the other hand, EPHA2 regulates CSCs along with other cancer properties. It is shown to regulate CSCs of glioblastoma (22), non-small cell lung carcinoma (23), nasopharyngeal carcinoma (24) and oral squamous cell carcinoma (OSCC) (25). The majority of the reports of EPHA2 regulating CSCs show the ligand-independent phosphorylation on S897 to be a critical regulator of the property (24, 26, 27). Another recent report has shown that EPHA2 leads to the transcriptional activation of KLF4 through ERK activation (25). Even though the role of EPHA2 receptor in CSCs is well established, the mode of activation of EPHA2 or the ligand molecule that interacts with it in CSCs, leading to these biological effects, is not extensively studied.

Ephrin-B1 is a ligand typically shown to bind with EphB1 or EphB2 receptor, and is well-explored in the context of stem cells. Ephrin-B1-mediated EphB signaling is shown to be important for the proliferation of dental pulp stem cells, and tissue repair (28, 29). Similarly, the neural progenitor compartment is also regulated by this signaling (30). The role of this signaling is extensively studied in mouse intestinal crypts, where the reverse gradient of EphB2/B3 and Ephrin-B1 expression and signaling thereby regulates the stem cell compartment position, proliferation, and differentiation (7). In spite of its well-established role in normal stem cell regulation, it’s contribution to cancer stem cells (CSCs) and cancer is poorly explored. However, a correlative analysis has shown that Ephrin-B1 is a marker for the aggressiveness of bladder cancer (31). Recently, we have reported that ablation of TUBB4B diminishes the self-renewal ability of oral CSCs *in vitro*, by reducing the surface localization of Ephrin-B1, implicating its role in oral cancer (32).

The current study was initiated with the investigation of the signaling pathways, their ligands and receptors, regulating self-renewal of oral cancer cells. Since our results pointed out the importance of Ephrin-B1 and the signaling mediated by this ligand, we explored the role of this signaling in regulating the self-renewal ability of oral cancer cells, which leads to recurrence and poor prognosis. Here, we present evidence for the interaction of Ephrin-B1 with EPHA2 in the same cancer cells, leading to the phosphorylations of Ephrin-B1. Further, we explored the possibility of this Ephrin-B1-EPHA2 interaction in the regulation of oral cancer prognosis using mouse models and OSCC primary samples.

## Results

### Eph/Ephrin signaling is regulated in parallel to the enrichment of self-renewing cancer cells in oral cancer

Since self-renewing cancer cells are the culprits of cancer properties like tumor initiation, metastasis, and recurrence, we explored the pathways regulating self-renewal ability in oral cancer cells. The surface proteins, usually the receptors and ligands of different pathways, critically regulate the properties of cancer cells. Sphere culture is an established way of enriching the self-renewing population, which we confirmed with sustained sphere formation efficiency up to three passages, as well as increased soft agar colony formation, increased expression of stem cell markers like *ALDH1A1*, *BMI1, KLF4, NANOG,* and *OCT4A* in sphere culture compared to monolayer cells (S1A-S1E Fig). To identify the surface proteins and pathways of self-renewal in oral cancer, we performed a proteomic analysis of the surface proteins and a phosphoproteomic analysis, using sphere culture in comparison to monolayer cells after SILAC labelling (S2A-S2B Fig). We identified 148 membrane proteins specifically enriched in sphere culture (Fig 1A, S1 Table). Up-regulation of some of the surface proteins obtained from the analysis was confirmed using western blotting of the membrane preparation (Fig 1B-1C). The surface localizations of a few molecules (TUBB, TUBA1C, TUBB2A, DSG3, TUBB4B, EFNB1) were confirmed using immunofluorescence and FACS analysis (S3A Fig, Fig 1D-1E). One of the molecules found to be enriched in spheres was Ephrin-B1 (EFNB1), which is a membrane-bound ligand involved in Eph/Ephrin signaling.

**Fig 1.** Eph/Ephrin signaling is regulated in parallel to the enrichment of self-renewing cancer cells in oral cancer. (A) SILAC-labelled HSC-4 oral cancer cells were grown in monolayer and sphere culture for 6 days (3D-spheres). Then we did membrane preparation followed by a SILAC based proteomic analysis using an LTQ–OrbitrapVelos ETD mass spectrometer. The Venn diagram shows the number of surface proteins enriched in spheres (148), monolayer (119), and common in both (285). (B) Some of the molecules obtained from proteomic analysis were confirmed in western blotting using lysates of 6-Day-old 3D-spheres (Sph) and monolayer (ML) of HSC-4 cells. (C) For ML and Sph, we normalized the expression with β-Actin. Then the fold change was calculated by dividing the relative expression in sphere by monolayer, and was plotted as mean ± SD values from 3 biological replicates. (D-E) HSC-4 Cells were grown in monolayer and sphere conditions for 3 days, and were used for FACS analysis to check surface expression of some of the molecules under non-permeabilizing condition. The percentage of cells showing surface expression of the indicated molecules in monolayer and spheres was plotted as mean ± SD values from 3 replicates. (F) As explained before, SILAC-labelled HSC-4 oral cancer cells grown as described earlier were used for phosphoproteomic analysis after enrichment of phosphopeptides using TiO_2_ beads. The pathways differentially regulated in spheres in the two independent experiments are shown in the Venn diagram. We identified 110 unique pathways differentially regulated in spheres. (G) Among the 110 unique pathways identified, the top 20 pathways involved in self-renewal are represented in the Pie chart. (H) HSC-4 cells were grown as monolayer or sphere for 3 days, and were used for FACS analysis for surface expression of the indicated molecules under non-permeabilizing condition. The percentage of cells showing surface expression of the indicated molecules in monolayer and spheres was plotted as mean ± SD values from 3 replicates. (I) Western blot analysis of the whole cell lysates of 3-Day-old Sph or ML of HSC-4 cells. (J) Data plotted as described before. Statistical analysis was done by an unpaired-two-tailed students t-test. *, **, ***, **** represents p<0.05, p<0.01, p<0.001 and p<0.0001, respectively.

Further, to identify critical pathways, we did two independent phosphoproteomic analyses using oral cancer cells, HSC-4. The peptides and proteins identified in replicates 1 and 2 are given in S2-S4 Tables. Our duplicate analysis identified 110 unique pathways differentially regulated in spheres compared to monolayer (Fig 1F, S5 Table). From the 110 pathways, we selected 20 pathways that are shown to be linked to self-renewal either in normal or cancer contexts, are given in S6 Table and Fig 1G. Phosphorylations of some of the molecules were confirmed using western blotting (S3B Fig).

Supporting the observation of surface enrichment of Ephrin-B1 in our surface proteomic analysis, we observed that Eph/Ephrin signaling is one of the signaling pathways differentially regulated in the sphere condition. Out of the 17 molecules involved in this pathway that were differentially phosphorylated upon sphere condition, the ligands and receptors involved were EPHA2, EPHB2, and Ephrin-B1 (S7 Table). While the ligand Ephrin-B1 and receptor EPHA2 showed hyperphosphorylation, EPHB2 exhibited hypophosphorylation upon sphere formation (S7 Table, S3C Fig). The results suggested that self-renewal enrichment is positively correlated to Ephrin-B1 and EPHA2, whereas it showed a negative correlation with EPHB2. A FACS analysis was performed to confirm the surface expression of receptors and ligands, which may lead to signaling. In accordance with the phosphorylations observed, the surface expression of Ephrin-B1 and EPHA2 increased with a concomitant decrease in EPHB2 surface expression in the sphere condition (Fig 1H, S3D Fig). To further explore the significance of this observation, we took 4 oral cancer cell lines to analyze the differential expression in spheres and monolayer cells by western blot. Western blots revealed that EPHA2 receptor and Ephrin-B1 ligand are enriched in spheres across all the cell lines, while EPHB2 receptor showed no significant change in the total expression except in HSC-4 cells, where it showed a significant reduction (Fig 1I-J). To confirm the specificity of bands of EPHA2 and Ephrin-B1 in the western blot, we used Lentiviral shRNA-mediated knock-down of the ligand and the receptor molecules. Consistent with the 57% reduction of *EPHA2* transcript, we observed a 51% reduction of EPHA2 in the western blot for the 125kDa band upon knock-down of EPHA2 (S4A-S4C Fig). Likewise, there was a comparable reduction of Ephrin-B1 in the western blot for the 51kDa band when Ephrin-B1 was knocked down (33% for both transcript and protein) (S4D-S4F Fig). Taken together, our results show that there is an increased surface expression of the ligand Ephrin-B1 and the receptor EPHA2 upon the induction of self-renewal in oral cancer cells. Thus, the chance of interaction of EphrinB1 with EPHA2 is higher in spheres compared to monolayer.

### Ephrin-B1 engages with EPHA2, and leads to differential phosphorylation in the self-renewal-enriched sphere condition

As our results indicated that in sphere condition, Ephrin-B1 might interact with EPHA2, we performed a co-immunoprecipitation (Co-IP) experiment to check whether the ligand binds to EPHA2 or EPHB2. As expected from the surface expression pattern, the level of Ephrin-B1 and its interaction with EPHA2 was higher in spheres compared to monolayer cells (Fig 2A). At the same time, Ephrin-B1 did not interact with EPHB2 in sphere or monolayer (Fig 2A). Then, the interaction of Ephrin-B1 ligand with EPHA2 receptor was validated using reverse Co-IP (Fig 2B).

**Fig 2.** Ephrin-B1 engages with EPHA2, and leads to differential phosphorylation in the self-renewal enriched sphere condition. (A) HSC-4 cells were grown as monolayer or 3-Day-old spheres were used for the Co-IP experiments. Co-IP was performed with mouse Anti-Ephrin-B1 or mouse IgG (3µg) along with 7% input. The eluate was probed for the indicated molecules. (B) A reverse Co-IP experiment was performed with Rabbit Anti-EPHA2 or Rabbit IgG along with a bead only control, and 7% input. (C-D) western blotting in the whole cell lysate of monolayer or 3-Day-old spheres to check phosphorylation of indicated molecules. (E) For ML and Sph, we normalized the expression with the total form. Then the fold change was calculated by dividing the relative expression in sphere by monolayer, and was plotted as mean ± SD values from 3 biological replicates. (F) Western blotting in the whole cell lysate of 3-Day-old 3D-spheres to check phosphorylation of indicated molecules in single knock-down cells of EPHA2 or Ephrin-B1 alone or in a 1:1 mixture of both single knock-down cells. (G) We normalized the expression with the total form. Then the fold change was calculated by dividing the relative expression in knock-down by control, and was plotted as mean ± SD values from 3 biological replicates. (H) Western blotting in the whole cell lysate of 3-Day-old 3D-spheres to check phosphorylation of indicated molecules in HSC-4 cells overexpressed with plasmid control (Ephrin-B1 RFP), Ephrin-B1 Y324F RFP (We used plasmid containing Ephrin-B1 sequence of mouse. Y323 residue of mouse Ephrin-B1 protein corresponds to Y324 in human protein, so mentioned as Y324 throughout the manuscript), and Ephrin-B1 Y317F RFP (Y316 residue of mouse Ephrin-B1 protein corresponds to Y317 in human protein, so mentioned as Y317 throughout the manuscript). (I) We normalized the expression with the total form. Then the fold change was calculated by dividing the relative expression in the mutant by the wild type control, and was plotted as mean ± SD values from 3 biological replicates. (J) *In vitro* kinase assay was performed with EPHA2 kinase obtained from the eluate of the co-immunoprecipitation with either Anti-EPHA2, IgG or bead-only control, and a small stretch of synthesized peptides with specific tyrosine residues, as mentioned, acted as acceptor for the kinase assay. Negative control for the kinase assay lacked the kinase (IP eluate). The graph represents the IP eluate dose curve from duplicate experiments. Statistical analysis was done by an unpaired-two-tailed students t-test. *, **, ***, **** represents p<0.05, p<0.01, p<0.001 and p<0.0001, respectively.

Next, to check whether the interaction of Ephrin-B1 ligand with EPHA2 receptor leads to phosphorylation, we performed western blotting experiment to identify their phosphorylation, on residues that are reported to be involved in biological activities (Fig 2C-2E). EPHA2 residues, Serine 897 (S897), Tyrosine 772 (Y772), and Tyrosine 594 (Y594) were showing hyper-phosphorylation in spheres. Similarly, Ephrin-B1 residues Y324/329 and Y317 were exhibiting hyper-phosphorylation in spheres. Contrary to these hyper-phosphorylations, Y588 of EPHA2 was showing reduced phosphorylation in spheres. As we expected, phosphorylation of EPHB2 at Y594/604 residue was showing reduced phosphorylation in spheres, which implies that EPHB2 activation may not be important in spheres (Fig 2C-2E). So our results show that when there is an enrichment of self-renewing population, there is an increased expression of Ephrin-B1 and EPHA2, leading to increased phosphorylations of EPHA2 at S897, Y772, Y594, and Ephrin-B1 at Y324/329 and Y317.

The increased phosphorylation we observed could be due to the EPHA2-Ephrin-B1 interaction, or it may be completely independent of it. To check that, we evaluated the change in the phosphorylation pattern when the ligand or receptor is depleted. Here, we checked how much of the receptor phosphorylation is blocked when the ligand is reduced and *vice versa*. None of the phosphorylations of EPHA2 that were specifically upregulated in spheres (S897, Y594 and Y772) were reduced upon Ephrin-B1 knock-down, suggesting that these phosphorylations are not exclusive to Ephrin-B1 interaction. At the same time, the phosphorylations of Ephrin-B1 at Y324/329 and Y317 were reduced significantly upon EPHA2 knock-down, which suggested that these Ephrin-B1 phosphorylations are EPHA2-dependent (Fig 2F-2G). Since we focused on these EPHA2-dependent Ephrin-B1 phosphorylations, we confirmed the authenticity of the bands by different methods. As shown in the full blot, the phosphorylation that was reduced upon the knock-down of EPHA2 was slightly heavier than the native form, and was around 55kDa (S5A-S5B Fig). Subsequently, we performed cloning to mutate these specific tyrosine residues at 324 or 317 to phenylalanine in the Ephrin-B1 RFP plasmid to validate the phosphorylations (S6A-S6B Fig). HSC-4 cells overexpressing these mutants were used to check the phosphorylations by western blot. Ephrin-B1 Y324F and Ephrin-B1-Y317F mutants showed a significant reduction in the phosphorylation of the corresponding residues (Fig 2H-2I). Additionally, EPHA2-dependent Ephrin-B1 phosphorylations were confirmed using an *in vitro* kinase assay where the Ephrin-B1 peptides containing Y324 and Y329 or Y317 residues were used as acceptor, and the Co-immunoprecipitated EPHA2 from spheres of oral cancer cells was used as the kinase. We show that Ephrin-B1 peptides are phosphorylated by EPHA2 in a dose-dependent manner, which was validated with IgG and Bead only control to negate non-specificity of antibody during the Co-immunoprecipitation experiment. A reaction lacking IP eluate served as the negative control for the kinase reaction. A peptide containing Y588/Y594, representing the autophosphorylation residues of EPHA2, was used as the positive control for the experiment (Fig 2J).

To visualize this EPHA2/Ephrin-B1 interaction leading to phosphorylation of Ephrin-B1, we exploited the proximity ligation assay (PLA). When we used EPHA2 and pEphrin-B Y324/329 primary antibody combinations for this assay in oral cancer sphere cells, we observed EPHA2/Ephrin-B Y324/329 interaction as red PLA foci, where the majority of them were on the cell membrane. While there were about 2 foci on average on cells in the test dish, the PLA dots observed on the secondary antibody control and the IgG control were negligible (Fig 3A-3D). In conclusion, our biochemical assays confirmed that the interaction of Ephrin-B1 with EPHA2 on the cell membrane leads to the phosphorylation of Ephrin-B1 at Y324/329 and Y317, while it is unclear whether it leads to the phosphorylation of any residues of EPHA2.

**Fig 3.** Ephrin-B1 engages with EPHA2 at the cell membrane, and leads to Ephrin-B1 signaling in the self-renewal enriched sphere condition. HSC-4 cells grown as 3 Day-old spheres were used for the experiments. (A) Proximity ligation Assay (PLA) performed with mouse EPHA2 and rabbit pEphrin-B1 Y324/329 antibodies. Red spot indicates PLA signal. The cell membrane was marked using Wheat Germ Agglutinin (WGA)- Alexa Fluor 488. The white arrows show the membrane localization of the PLA loci. (B) Zoomed-in images of the rectangle shown in A. (C) PLA of secondary control (without primary antibodies) and IgG control containing rabbit and mouse IgG antibodies. (D) Quantification of PLA foci per cell in 4 fields of 60X images. 37, 29, and 45 cells were counted for secondary control, IgG control, and the test, respectively. Statistical analysis was done by an unpaired-two-tailed students t-test. **** represents p<0.05, p<0.01, p<0.001 and p<0.0001, respectively.

### EPHA2/Ephrin-B1 interaction supports the self-renewing cancer stem cell (CSC) pool in oral cancer cells

Since we anticipated that EPHA2/Ephrin-B1 interaction supports CSCs in oral cancer, we evaluated the change in CSC pool with the depletion of the receptor EPHA2 and the ligand Ephrin-B1. For this, we used a published CSC reporter, ALDH1A1-DsRed2 (32, 33). It was made by cloning the ALDH1A1 promoter upstream of DsRed2 in the DsRed2N1 plasmid (Fig 4A). Lentiviral shRNA-mediated knock-down was performed in the reporter cell lines (Fig 4B-C). We analyzed ALDH1A1-positive population in the reporter cells with knock-down of EPHA2 or Ephrin-B1 through FACS analysis. Our results showed that knocked-down cells in the sphere condition had a higher reduction in ALDH1A1-positive population compared to CSC-enriched monolayer (Fig 4D-G). Both EPHA2 or Ephrin-B1-depleted cells exhibited a significant reduction in the ALDH1A1-positive CSC pool compared to control cells in the spheres as well as CSC-enriched monolayers (Fig 4E-F). Further, we analyzed the ALDH1A1-positive population in dual knock-down cells (both receptor and ligand), which showed superior reduction in ALDH1A1-positive CSC pool compared to single knock-down cells (Fig 4G). It suggests that EPHA2/Ephrin-B1 interaction is important for the maintenance of ALDH1A1-positive CSC pool.

**Fig 4.** EPHA2/Ephrin-B1 interaction regulates self-renewing cancer stem cell pool in oral cancer cells. (A) The vector construct of ALDH1A1-DsRed2 reporter. It was made by cloning the ALDH1A1 promoter gene upstream of DsRed2 in the DsRed2N1 plasmid. (B) HSC-4 Reporter cells were made by stable selection with G418 and followed by FACS sorting of the ALDH1A1-DsRed2 positive cells. (B-C) Western blot to check the knock-down of the indicated molecules in HSC-4 ALDH1A1-DsRed2 cells, and the data represent fold change based on the control cells, after normalizing with β-Actin. (D-E) FACS analysis to check ALDH1A1 population in knock-down cells in 3-Day-old monolayer or spheres. (E) Data indicate the percentage of ALDH1A1-positive CSC population, plotted as mean ± SD values from 4 replicates. Statistical analysis was done by an unpaired-two-tailed students t-test. *, **, ***, **** represents p<0.05, p<0.01, p<0.001, and p<0.0001 respectively.

To corroborate the results obtained by the CSC reporter cells, we performed more robust self-renewal functional assays, like extreme limiting dilution assay (ELDA) (34). For that, we used single knock-down of either receptor or ligand and dual knock-down of both for ELDA analysis (S7A-S7B Fig). First, we did an *in vitro* serial dilution sphere formation assay to evaluate the effect of the knock-down of the ligand and the receptor (Fig 5A). The depletion of EPHA2 significantly reduced the CSC frequency in oral cancer cells (4.5-fold reduction in RCB1015 and 2.5-fold reduction in HSC-4 cells (Fig 5B-5C, S7C Fig). Similarly, knock-down of Ephrin-B1 caused a 2.5-fold reduction in the CSC pool (Fig 5B-5C). Although we observed a significant reduction in the CSC frequency, the single knock-down of either the receptor or ligand did not substantially change the size of the spheres or the number of spheres formed in each dilution (Fig 5A, S7C-S7D Fig). Interestingly, the dual knock-down of EPHA2 and Ephrin-B1 drastically reduced the CSC frequency (6-fold reduction), sphere size and number of spheres formed in each dilution (Fig 5A-5C, S7D-S7E Fig). Thus, our observations show that abolishing both ligand and receptor in the same cell, disrupting the EPHA2/Ephrin-B1 interaction, drastically depletes CSCs. Then, to analyze whether *in vitro* studies are concordant with the *in vivo* scenario, we did an *in vivo* serial dilution xenograft assay. Similar to *in vitro* results, the CSC-frequency was significantly reduced upon depletion of EPHA2 or Ephrin-B1 (Fig 5D-5G, S8A-S8B Fig). Taken together, all these results show that abrogation of EPHA2/Ephrin-B1 interaction abolishes CSCs *in vitro* and *in vivo*. Furthermore, we did an *in vitro* serial dilution sphere formation assay with Y324F or Y317F mutants of Ephrin-B1 to check the dependency of self-renewal on the phosphorylation of these residues. The mutation of tyrosine 324 to phenylalanine of Ephrin-B1 (Ephrin-B1 Y324F) caused a 1.7-fold reduction in the CSC frequency in oral cancer cells (Fig 5H-5J, S8C Fig). Similarly, Thus, our results so far confirmed that the EPHA2-mediated phosphorylations of Ephrin-B1 on Y324 and Y317 are critical for the self-renewal of oral cancer cells.

**Fig 5.** Abrogation of EPHA2/Ephrin-B1 interaction depletes self-renewing cancer stem cell pool in oral cancer cells. (**A**) RCB1015 cells with knock-down of EPHA2, Ephrin-B1 or both molecules were seeded at different serial dilutions in Poly-HEMA-coated 96-well plate and were grown as spheres for 6 days. The image shows the spheres formed in the dilution of 10,000 cells/well. The scale bar represented 100µm. (B) Cancer stem cell frequency calculated by ELDA (C) The graph represents a log fraction plot obtained from ELDA analysis, and slope of it represents CSC frequency. (D) For the serial dilution xenograft formation assay, cells were serially diluted and were injected subcutaneously on the flanks of NSG mice. The first two figures in D represent the image of tumors collected after 14 days from 10^6^, and 10^5^ dilutions respectively while tumors were collected after 40 days from 10^4^ dilution. (E) Representative images of tumors collected after 17 days from all the dilutions. (F-G) Tables show a summary of ELDA analysis, and graphs represent the log fraction plot obtained from ELDA analysis. (H) 6-Day spheres formed of HSC-4 cells overexpressed with Ephrin-B1 RFP or Ephrin-B1 Y324F RFP in a dilution of 1000 cells/well of respective cells. (I-J) The table shows a summary of ELDA analysis, and the graph represents a log fraction plot obtained from ELDA analysis. (K) 6-Day spheres formed of HSC-4 cells overexpressing respective constructs as described before. (L-M) The table shows a summary of ELDA analysis, and the graph represents a log fraction plot obtained from ELDA analysis.

### *Cis*-interaction of EPHA2 and Ephrin-B1 regulates the self-renewing cancer stem cell (CSC) pool in oral cancer cells

Since Eph receptors and Ephrin ligands are reported to interact in *cis* (within the same cell) as well as *trans* (between neighboring cells), we evaluated these possbilities in our system. In our earlier assays of self-renewal, we have used dual knock-down system, where the receptor and the ligand of the same cell are depleted. So, in a dual knock-down system, both *cis* and *trans* interactions are compromised. Then we used the single knock-down cells, where both interactions are weakened. At the same time, when we mix these cells in equal proportions, they can restore the *trans* interaction because EPHA2 knock-down cells will express Ephrin-B1, and Ephrin-B1-depleted cells will have EPHA2. Mixing of the single knock-down cells failed to rescue the phosphorylation of both the residues of Ephrin-B1, implying that the interaction is a *cis*-interaction (Fig 2F). Then, to determine whether this *cis*-interaction leads to phosphorylation of Ephrin-B1 at tyrosine 324, we analyzed multiple confocal images of the PLA experiment. It showed a higher number of PLA foci indicating EPHA2/Ephrin-B1 Y324 interactions were on the cell membranes without cell contact than on the cell-cell junctions, which suggested *cis*-interaction leading to the phosphorylation of Ephrin-B1 in spheres (Fig 6A-6B).

**Fig 6.** EPHA2/Ephrin-B1 *cis-*interaction regulates self-renewal. (A) Images of PLA showing the foci at isolated cell membranes without cell-cell contact (*cis,* arrow marks), or cell-cell junctions where we cannot distinguish whether it is *cis* or *trans*. (B) The distribution of PLA foci/cell from 63 cells in 4 fields of 60X images. Statistical analysis was done by an unpaired-two-tailed students t-test. **** represents p<0.0001. (C) Diagrammatic representation of FRET facilitated photoswitching used in our experiment. (i) mT-Sapphire-C1 was excited with 405 nm and emission collected with 480/40 nm filter. For PSmOrange-C1, the excitation was with 560 nm and emission collected with 575/30 nm filter. (ii) For photoswitching of PSmOrange-C1, a 488 nm laser was used. The far-red form of PSmOrange-C1 was excited with 639 nm, and emission was collected with 650/25 nm. (iii) When mT-Sapphire-C1 is excited with 405 nm, it emits light in the range of 500-570 nm, which induces photoswitching of PSmOrange-C1 if it is within 10 nm. The photoswitching can be confirmed with an excitation of 639 nm and emission of 650/25 nm. (D) HSC-4 cells were transiently transfected with PSmOrange C1-EPHA2 and mT-Sapphire C1-Ephrin-B1 plasmids. Then, these cells were grown as spheres for 3 days. Then, we imaged FRET facilitated photoswitching in live cells where we performed a sequential acquisition, as Ex. 639 nm/ Em. 650/25 nm, Ex. 560 nm/ Em. 575/30 nm, Ex. 405 nm/ Em. 488/40 nm. Further, a pulse of 30 iterations of excitation of 405 nm was given. Then, the next cycle was Ex. 639 nm/ Em. 650/25 nm, Ex. 560 nm/ Em. 575/30 nm (E-F) Table represents the summary of ELDA analysis, and the graph represents a log fraction plot obtained from ELDA analysis of mixed cells containing an equal proportion of EPHA2 knock-down cells and Ephrin-B1 knock-down cells, and the slope of it represents CSC frequency.

To ascertain the *cis*-interaction of EPHA2 and Ephrin-B1 in live oral cancer cells in a self-renewing sphere condition, we performed a FRET facilitated photoswitching with PSmOrange-C1 and mT-Sapphire-C1, which are established photoswitching pairs (35–37). mT-Sapphire C1 excites at 399 nm (ranges from 390-420 nm) and emits a blue-green light at 511 nm (ranges from 500-570 nm) while PSmOrange-C1, which is initially orange (excitation-548 nm, emission-565 nm) undergoes photoswitching upon irradiation with blue-green light to a far red form (excitation-636 nm, emission-662 nm) (Fig 6C). For this experiment, we made PSmOrange-C1-EPHA2 and mT-Sapphire-C1-Ephrin-B1 clones by amplifying EPHA2 or Ephrin-B1 from corresponding plasmids, followed by restriction digestion and ligation into PSmOrange-C1 and mT-Sapphire-C1 plasmids (S9A-S9B Fig). Further, we transiently transfected them into oral cancer cells. If EPHA2 and Ephrin-B1 are in proximity of less than 10 nm, the emission wavelength of mT-sapphire causes photoswitching of PSmOrange to a far red variant.

PSmOrange C1-only control assured photoswitching property of PSmOrange and was the positive control for the experiment (S10A Fig), while the combination of empty vectors of PSmOrange-C1 and mT-Sapphire was kept as the negative control (S10B Fig). For our analysis, we selected cells co-expressing both constructs, which are not in proximity. We observed photoswitching on the membrane of the same cell co-expressing both PSmOrange-C1-EPHA2 and mT-Sapphire-C1-Ephrin-B1, which confirmed the *cis*-interaction of EPHA2 and Ephrin-B1 (Fig 6D, S10C Fig). Next, to check whether the *cis* interaction is involved in self-renewal, we performed an *in vitro* serial dilution sphere formation assay with a mix of equal proportions of EPHA2 knocked-down cells and Ephrin-B1 knocked-down cells. These mixed cells exhibited a 2.3-fold reduction in the CSC frequency in oral cancer cells, which confirms the role of *cis*-interaction in self-renewal of CSCs (Fig 6E-6F, S11 Fig). Collectively, we show that Ephrin-B1 ligand interacts with EPHA2 receptor in spheres in a *cis*-manner, and leads to the phosphorylations of residues at Y324/329 and Y317 of Ephrin-B1 ligand, which contributes to the self-renewal ability of CSCs.

### Abrogation of EPHA2/Ephrin-B1 interaction increases the overall survival in a mouse model of oral cancer

Since CSCs exhibiting self-renewal ability are shown to critically regulate the prognosis of cancer, we evaluated the role of this signaling pathway in the overall survival using an orthotopic mouse model of oral cancer. When mice injected with EPHA2 or Ephrin-B1 knocked-down cells exhibited a delayed tumor progression, mice injected with dual knocked-down cells displayed a drastic retardation in tumor progression (Fig 7A-7B). Then, we used Kaplan-Meier analysis to study the prognosis. Though, there was a trend in the Ephrin-B1-depleted group to show improved overall survival, single knock-down of either receptor or ligand was incapable of significantly altering the overall survival of the mice. In contrast, mice injected with dual knocked-down cells showed a significant improvement in overall survival (Fig 7C). Furthermore, the expression of self-renewal molecules like SOX2 and OCT4A showed a reduction in the ligand or receptor-knocked-down tumors (S12A-S12B Fig). In agreement with this, xenograft tissue sections with knock-down of receptor or ligand (S12C-S12E Fig) showed reduced expression of ALDH1A1 (S12C, S12F Fig). To confirm the role of Ephrin-B1 phosphorylations in self-renewing CSCs *in vivo*, we used the serial sections of the orthotopic tumors for immunofluorescence. As expected, phosphorylation of Ephrin-B Y324/329 on the cell membrane was high in the regions where ALDH1A1 was high, compared to the ALDH1A1^low^ region (Fig 7D-7E). More important observation was the high co-localization of EPHA2 and phospho-Ephrin-B Y324/329 on the cell membrane of cells in the ALDH^high^ region as revealed by Pearson’s correlation analysis Scatter Plot (Fig 7D) and the Pearson’s correlation coefficient (Fig 7F). Our results clearly demonstrate the increased interaction of EPHA2 with Ephrin-B, thereby increasing the phosphorylation of Ephrin-B Y324/329 on the ALDH^high^ region. To summarize, the interaction of EPHA2 and Ephrin-B1 in the CSC niche supports the sustenance of CSCs, and the loss of EPHA2 and Ephrin-B1 in the same cell reduces the tumor progression and improves the prognosis by enhancing overall survival in mouse models.

**Fig 7.** Role EPHA2/Ephrin-B1 interaction supports ALDH1A1 population leading to poor prognosis in oral cancer mouse model. Overall survival analysis was performed using an orthotopic floor of the mouth mice model. 0.5×10^6^ cells were injected into the floor of the mouth region superficial to the mylohyoid muscle. RCB1015 shControl, shEPHA2 and shEphrin-B1 cells were injected into SCID mice, while RCB1015 shControl and shDual, cells were injected into NSG mice. (A) Representative image of mice with a tumor after 24 days. (B) We measured tumor size using a digital Vernier caliper at regular intervals and plotted tumor volume against the number of Days. Tumor volume was calculated using the formula (Height*Height*width)/2. (C) Kaplan-Meier survival analysis was performed using GraphPad Prism to check overall survival. (D) Xenograft serial sections were co-stained for ALDH1A1/Ephrin-B Y324/329 or EPHA2/Ephrin-B Y324/329. We chose two regions, one with high expression of ALDH1A1 (ALDH1A1^high^) and the other with low expression of ALDH1A1 (ALDH1A1^low^), and the same regions were identified on the next serial section that were stained for EPHA2/Ephrin-B Y324/329 and represented with an image of high magnification. The Scatter plots represent Pearson analysis between EPHA2 and phospho-Ephrin-B Y324/329 of ALDH1A1 high and low regions in the whole field. (E) Mean intensity of Ephrin-B Y324/329 was calculated from the cell membranes of 75 cells each from 3 fields of ALDH1A1 high and low regions. (F) ROI was drawn for cell membranes of 75 cells from 3 fields and calculated Pearson’s correlation coefficient between EPHA2 and phospho-Ephrin-B Y324/329 in ALDH1A1 high and low regions. Statistical analysis was done by an unpaired-two-tailed students t-test. *, **, **** represents p<0.05, p<0.01, p<0.001 and p<0.0001, respectively.

### EPHA2/Ephrin-B1 interaction critically regulates the prognosis of oral cancer

Data from our mouse model, demonstrating the role of EPHA2/Ephrin-B1 interaction in the regulation of overall survival, prompted us to explore the role of this signaling in overall survival of oral cancer patients. First, we checked whether there were any differences in the expression of the ligand and the receptors identified in our study, using a TCGA dataset of oral cancers compared to the normal tissues. The transcript levels of Ephrin-B1 (*EFNB1*) and EPHB2 (*EPHB2*) were significantly upregulated in the malignant tissues compared to their normal counterparts. Though there was a trend of upregulation, the levels of EPHA2 (*EPHA2*) did not significantly change in the two conditions (S13A Fig). Yet, when we analyzed the correlation of Ephrin-B1 with the receptors using a hierarchical clustering heat map analysis of TCGA data, the correlation with *EPHA2* was higher than with *EPHB2* receptor (S13B-S13C Fig). Then, we evaluated the protein expression of EPHA2, EPHB2, and Ephrin-B1 in relation to ALDH1A1 on the primary oral cancer tissue sections to check the dependency of this signaling on CSCs. An immunohistochemical analysis of oral cancer samples co-stained with these four molecules revealed that while the expression of EPHA2 receptor and Ephrin-B1 ligand is confined to certain clusters, the expression of EPHB2 is uniform throughout the section (Fig 8A). More specifically, some regions of the tissue sections had high expression of both EPHA2 and Ephrin-B1, which showed a better co-localization than the Ephrin-B1-EPHB2 co-localization, as revealed by Pearson’s correlation analysis (Fig 8B). Hence, in accordance with our *in vitro* cell line data, the primary oral cancer sample data also suggest EPHA2-Ephrin-B1 co-localization leads to signaling within the region positive for ALDH1A1, defining the CSC niches of the tumors. The expressions of both Ephrin-B1 and EPHA2 were observed in the same cell, suggesting the *cis*-interaction, which we deduced from our biochemical analysis. Here, we observed that the ALDH1A1 population is restricted to the region where EPHA2-Ephrin-B1 co-localization is seen (Fig 8A), supporting the notion that the interaction of EPHA2 and Ephrin-B1 in the same cell enriches self-renewing CSCs in oral cancer.

**Fig 8.** EPHA2/Ephrin-B1 interaction in oral cancer tissues. (A) OSCC patient samples were co-stained for the indicated molecules using tagged primary antibodies, EPHB2-Alexa 405, EPHA2-Alexa 488, EphrinB1-Alexa 680, and ALDH1A1-Alexa 568. (B) From multiple images at 40X, we chose 12 ROI to check the co-localization using Pearson’s correlation analysis. **, ***, **** represents p<0.01, p<0.001 and p<0.0001, respectively.

Next, we tested whether the expression of *EFNB1* and *EPHA2* correlates with overall survival. Although patients with the high expression of either the ligand or receptor did not show any difference in the prognosis, the high expression of both the receptor and ligand exhibited poor overall survival compared to the group showing the low expression of both (Fig 9A-9C). Since the prognosis of oral cancer is dependent on the recurrence of the disease, we also studied that aspect. We segregated oral cancer patients into the recurrent and disease-free category, with a cut-off of 15 months’ disease-free survival. Then a tissue microarray immunohistochemical (IHC) analysis was performed on recurrent and disease-free OSCC samples using EPHA2 and Ephrin-B1 antibodies (Fig 9D). Irrespective of the survival status, the expression of the receptor was high in all tumor tissues. On the other hand, Ephrin-B1 expression was limited to the outer layers of tumor bundles rather than the core of the bundle. Then, tumor bundles in the samples were scored based on the co-expression of high EPHA2 (EPHA2^Hi^) with concomitant Ephrin-B1 positivity. Then, the percentage of tumor bundles with co-localization of EPHA2^Hi^ and Ephrin-B1^+^ in each sample was calculated. The co-expression of the ligand Ephrin-B1 and the receptor EPHA2 offers a chance for their interaction to modulate signaling. The recurrent group showed a higher percentage of tumor bundles (>30%) with co-localization of EPHA2^Hi^ and Ephrin-B1^+^ than the disease-free group (Fig 9E).

**Fig 9.** Role of EPHA2/Ephrin-B1 interaction in the prognosis of oral cancer patients. (A-C) A TCGA data set was used to study overall survival in HNSCC patients. Patients were categorized according to the level of expression of EPHA2 and Ephrin-B1, as described under Materials and Methods. The table represents the summary of statistics (D) A tissue microarray IHC was performed on serial sections of recurrent or disease-free samples Images from three patients from each group are shown (E) Then, tumor bundles in the samples were scored based on the co-localization of high expression of EPHA2 (EPHA2^Hi^) and Ephrin-B1 in the sample. Then, the percentage of tumor bundles with co-expression of EPHA2^Hi^ and Ephrin-B1^+^ in each sample was calculated. (F) The patients were segregated into two groups, using a cut-off of 30% of tumor bundles with co-expression of EPHA2^Hi^ and Ephrin-B1. A Kaplan-Meier survival graph for disease-free survival was plotted between those two groups.

Kaplan-Meier survival analysis showed that patients with a lower extent of co-expression of Ephrin-B1^+^ and EPHA2^Hi^ (<30% of the tumor bundles) show better disease-free survival compared to patients with a higher extent of their co-expression (>30%) (Fig 9F). Thus, our IHC analysis of the tissue microarray reinforced our hypothesis that the interaction of Ephrin-B1 with EPHA2, in *cis*, critically regulates recurrence of oral cancer.

## Discussion

Recurrence is a major driving factor for poor prognosis in many cancers, including lung, breast, bladder, and oral cancer (38–42). Among these cancers, oral cancer is a major burden in Asian countries, especially in India. Recurrence occurring as loco-regional or distant metastasis leads to the death of one out of two oral cancer patients (42, 43). The recurrence depends more on the self-renewal property of cancer cells than on the stage and grade of the cancer (44, 45). Therefore, it is essential to target these self-renewing cancer stem cells (CSCs) for which we need to identify surface proteins and pathways that regulate them. In self-renewal-enriched spheres, we identified the enrichment of the surface expression of Ephrin-B1 ligand, which is a mediator of Eph-Ephrin signaling. Moreover, a phosphoproteomic analysis identified Eph-Ephrin signaling as one of the differentially regulated pathways in spheres. The phosphorylation pattern and enrichment of the ligand and receptors in the sphere condition suggested the possibility of Ephrin-B1/EPHA2 signaling, supporting CSCs (Fig 1). Though cross-signaling is possible for Eph-Ephrin signaling, the majority of studies have shown that EPHA2 interacts with Ephrin-A, while Ephrin-B1 interacts with EPHB. In this context, if we find that EPHA2 interacts with Ephrin-B1 to support CSCs, it will be a novel interaction that can be targeted for eradicating CSCs.

Though Ephrin-B1 is established to regulate normal stem cells, its role in CSCs is less investigated except for the first report from our lab, which showed that the regulation of surface expression of Ephrin-B1 by TUBB4B is essential for the support of CSCs in oral cancer (32). Ephrin-B1, however, is reported to be overexpressed in many cancers, including oral cancer (32, 46, 47). Also, the over-expression is correlated to angiogenesis, aggressiveness of tumor, and recurrence, though the molecular mechanism by which it regulates cancer progression is elusive (46). Even though the EPHB2 receptor is reported as the regular binding partner for Ephrin-B1 in different contexts, as stem cell-homeostasis of intestinal crypt as well as cancer. During cancer progression, EPHB2/Ephrin-B1 signaling regulates invasion, metastasis, and immunomodulation (7, 18, 48). But whether the same receptor molecule interacts with Ephrin-B1 in the CSC context is unknown. Since our findings suggested that Ephrin-B1 and its signaling are crucial in regulating the self-renewal of cancer cells, it is essential to identify the modulators of Ephrin-B1 to regulate self-renewal.

The receptor molecules obtained from our phospho-proteomic analysis, EPHB2 and EPHA2, are reported to be overexpressed in HNSCC and oral cancer, respectively (25, 49). Higher expression of these receptor molecules in cancer patients leads to poor prognosis in patients, implying that they have a role in the progression of cancer (25, 50). Consistent with this, our data revealed that *EPHB2* transcripts and its protein are high in oral cancer, suggesting its role in the regulation of malignant properties (S12A Fig, Fig 8A). Although our results did not support the role of EPHB2 in the self-renewal in oral cancer, it is reported to enrich CSCs to impart sorafenib resistance in hepatocellular carcinoma, by the up-regulation of the WNT-β-catenin pathway (51). Despite that, its expression is a predictive marker for relapse-free survival in colorectal cancer, gastric cancer, and breast cancer, disproving its role as a universal positive regulator of CSCs and recurrence in all malignancies (52–54). Even if EPHB2 does not regulate the self-renewal property of CSCs, the signaling mediated by the receptor might have critical roles in other malignant properties. The EPHB-Ephrin-B interaction is reported to induce fibroblast remodeling to generate cancer-associated fibroblasts (CAFs), and increase invasion and metastasis (55).

As we consider the second receptor EPHA2, it is demonstrated to regulate CSCs of many cancers, including oral cancer, which aligns with our findings (22, 23, 25). Additionally, our study proposes the role of EPHA2 receptor in regulating recurrence, as we observed the co-expression of EPHA2 receptor and Ephrin-B1 ligand correlates with recurrence in oral cancer patients (Fig 9D-9F, S13A Fig). Since we observed that there is enhanced surface expression of Ephrin-B1 ligand and EPHA2 receptor when there is an induction of self-renewal (Fig 1I-1J), we investigated whether they interact to initiate signaling. Our co-immunoprecipitation experiments indicated that the co-expression of EPHA2 and Ephrin-B1 in the self-renewal-enriched sphere condition results in their interaction (Fig 2A-2B). Our western blot analyses showed that in the self-renewal-enriched condition, there is differential regulation of phosphorylation in the ligand as well as receptors (Fig 2C-2E). We detected a variation in receptor phosphorylations upon self-renewal enrichment. Phosphorylations of EphB receptor at Y594/604, and Y588 of EPHA2 were reduced, while S897, Y772, and Y594 residues of EPHA2 receptor were increased. When we used the single knock-down cells to evaluate the interdependence of Ephrin-B1 and EPHA2 for the differentially regulated phosphorylations, only the ligand phosphorylations were dependent on EPHA2, whereas receptor phosphorylations were independent of Ephrin-B1 interaction (Fig 2F-2G). The specificity of the residues of Ephrin-B1 phosphorylations was confirmed when we observed a reduction of phosphorylations of Y324/329 and Y317 upon overexpression of Y324F and Y317F mutants (Fig 2H-2I). Further, *in vitro* kinase assay substantiated that EPHA2 phosphorylates Ephrin-B1 residues Y324/329 and Y317 by (Fig 2J). Moreover, we could visualize the interaction of EPHA2 leading to the phosphorylation of Ephrin-B324/329 on the cell membrane of oral cancer cells under sphere culture using proximity ligation assay (PLA). The specific foci observed on the cell membrane reinforced our argument that EPHA2-Ephrin-B1 interactions could lead to the phosphorylation of Ephrin-B1 residues (Fig 3).

The functional consequences of the phosphorylations of ligands and receptors in the Eph/Ephrin family are not studied extensively. Yet, ligand-mediated phosphorylations of the receptors for the forward signaling are reported in different contexts. In many instances, the receptor phosphorylations are mediated by heterotypic receptors and other kinases. One such phosphorylation is of EPHA2 S897, a critical event regulating CSCs (24). This phosphorylation is shown to be ligand-independent, regulated by kinases other than EPH receptors, like Protein Kinase B (AKT), Protein Kinase C (PKC), and Ribosomal S6 kinase (RSK) (56–58). Another example of ligand-independent EPHA2 phosphorylation is at Y772, which is mediated by Focal Adhesion Kinases (59). Moreover, this ligand-independent EPHA2 Y772 phosphorylation is shown to promote proliferation and anchorage-independent growth (60). In accordance with these, we observed increased phosphorylation of EPHA2 S897 and Y772 in spheres that show anchorage-independent growth and enrichment of CSCs. Since these phosphorylations are independent of Ephrin ligands, they did not change with the loss of Ephrin-B1 (Fig 2J). Further, our results indicate that these heterotypic interactions of EPHA2 are independent of its *cis*-interaction with Ephrin-B1.

Though not reported in the CSC context, the classical ligand-dependent phosphorylations of EPHA2 are of Y588 and Y594, the two juxtamembrane domain residues that are reported to be autophosphorylated upon Ephrin-A1 ligand binding (1). These residues interact with the kinase domain in an unphosphorylated form, and this interaction is critical for the autoinhibition of the kinase domain (61). To the best of our knowledge, a direct substrate of EPHA2 kinase other than these two residues has not been reported. In our study, we found an increase in the phosphorylation of Y594 and a concomitant decrease in Y588 in sphere cells. So, our results suggest that the two autophosphorylations may not be interdependent. Crystal structure analysis has revealed that both Y588 and Y594 interact with the same pocket of the kinase domain (61). Nevertheless, whether these two residues are phosphorylated simultaneously, sequentially, or independently of each other has not been addressed so far. Upon phosphorylation, Y588 and Y594 bind to Vav2 and Vav3 guanine nucleotide exchange factors, respectively, to regulate Rac-mediated activities (62). The signaling events downstream of Vav3 may be synergistic or independent of Vav2 effects, depending on the context (63). In the sphere condition, where EPHA2 is interacting with Ephrin-B1, it leads to the phosphorylation of only Y594 of EPHA2, unlike when it interacts with Ephrin-A, its well-established ligand. The difference between these interactions and the subsequent phosphorylations of the receptor EPHA2 needs further evaluation. As we observed no change in the non-canonical activation of EPHA2 (Y772 and S897), it appears that *cis*-interaction of EPHA2 with Ephrin-B1 does not affect its interaction with other kinases. While we focused on phosphoforms with commercially available antibodies, we acknowledge the possibility of phosphorylation occurring at other EPHA2 residues, which we haven’t explored. In addition, the increased phosphorylation of EPHA2 may play a role in regulating CSCs, though this does not seem to rely on Ephrin-B1 interaction.

There are considerable reports arguing that the ligand-mediated receptor phosphorylations lead to forward signaling. However, reports of receptor-induced modifications of ligands leading to reverse signaling, are rare. Even if the functional significance of the Ephrin-B1 ligand phosphorylations, both Y324/329 and Y317, is not known, they are identified as phosphorylations induced by soluble EphB2 (64). The available reports do not reveal the interaction of Ephrin-B1 with receptors other than EphB. Since our focus was on Ephrin-B1-EPHA2 interaction, we further explored the signaling initiated by it. It is well established that Eph receptors and Ephrin ligands can interact in *trans* or *cis*, where conventionally the *trans*-interaction leads to signaling events, and the *cis*-interaction blocks the *trans*-interaction (14, 65). Interestingly, there is a very recent article that shows Ephrin-A1 fuels cervical cancer progression by its *cis*-interaction with EPHA2 (66). In agreement with this, here we show that EPHA2-Ephrin-B1 *cis*-interaction leads to signaling events. The pattern of PLA foci, especially the presence of foci on cell membranes where there is no cell-cell contact, suggested that the interaction could be a *cis*-interaction (Fig 6A-6B). This prompted us to explore the possibility in detail. As the co-culture of single knock-down of ligand or receptor retains the *trans* interaction while blocking the *cis* interaction, we performed our western blot analysis of the phosphoforms in that condition. Here, we showed that the *trans* interaction of the ligand and the receptor does not compensate for the loss of phosphorylation, suggesting that the *cis*-interaction of Ephrin-B1 with EPHA2 leads to Ephrin-B1 phosphorylation (Fig 2H-2I). To substantiate this novel finding we performed a live FRET facilitated photoswitching using PSmOrange-C1-EPHA2 and mT-Sapphire-C1-Ephrin-B1. We observed photoswitching in cells co-expressing both constructs, confirming the *cis*-interaction of the ligand and receptor (Fig 6C).

So far, our results demonstrate that in a self-renewal enriching condition, there is enhanced co-expression of EPHA2 and Ephrin-B1, increasing their interaction and phosphorylation. Although the ligands and receptors may interact in *cis* or *trans*, the interaction of Ephrin-B1 with EPHA2 in *cis* is a critical signaling event in the sphere condition. Next, we analyzed the functional significance of this novel EPHA2-Ephrin-B1 interaction and signaling. As our results indicated that the EphrinB1-EPHA2 interaction is important for self-renewal, we explored the relevance of this interaction in CSCs and recurrence. Our FACS analysis with ALDH1A1-DsRed2 CSC reporter cells showed the dependence of CSCs on EPHA2-Ephrin-B1 signaling. Our results showed a pronounced reduction in CSC frequency in dual knock-down cells compared to single knock-down cells, where the ligand-receptor interaction is blocked (Fig 4). Extreme limiting dilution assay (ELDA), a functional assay to evaluate the CSC frequency, corroborated the results of our reporter assay, both *in vitro* and *in vivo* (Fig 5, S7 Fig, S8A-B Fig). ELDA, using a mix of single knock-downs to restore *trans*-interaction, showed that reinstating the *trans*-interaction did not recover the loss of CSC pool (S10D Fig). Thus, it is evident that the *cis*-interaction is the major mode of CSC regulation. Next, to examine whether this regulation of CSCs by EPHA2-Ephrin-B1 *cis*-interaction decides the prognosis, we checked overall survival using orthotopic floor of the mouth mouse model (Fig 7A-7C). The drastic reduction we observed in the dual knock-down cells was indeed due to the depletion of CSCs as revealed by the depletion of ALDH1A1 and other self-renewal markers (S11 Fig). Further, analysis of the xenograft sections showed that the interaction of EPHA2 with phosphorylated Ephrin-B is significantly enhanced on cell membranes of CSC niches compared to other regions, supporting our claim that EPHA2-Ephrin-B1 *cis*-interaction is crucial to enrich CSCs (Fig 7D-7G). Next, we validated the clinical significance of our observations using a TCGA data set of HNSCC patients. Kaplan-Meier analysis showed that the patients with higher expression of both receptor and ligand had decreased overall survival time compared to patients having low expression of both, underscoring the significance of this interaction (Fig 9A-9C). Further, this observation was validated by IHC of the tissue microarray of OSCC samples. The pattern of expression of EPHA2 and Ephrin-B1 in OSCC samples suggest that the *cis*-interaction is clearly enhanced in malignant tissues of patients. More important observation was the significant increase of the EPHA2-Ephrin-B1 co-expression (and thereby their interaction) in patients who showed recurrence within 15 months compared to patients who showed disease-free-survival (Fig 9D-9F). To conclude, in the current study, we demonstrate the importance of Ephrin-B1 in the regulation of the self-renewal ability of oral cancer cells, which in turn regulates the chance of recurrence of the disease.

Though ligand molecules are implicated in cancer progression, the underlying mechanism is largely unknown. To the best of our knowledge, this is the first study revealing how Ephrin-B1 ligand regulates cancer progression and recurrence. The recent report (66) and our study challenge the conventional belief that receptors and their *trans*-forward signaling are the primary drivers of signaling, whereas *cis*-interactions serve only to block signaling. We argue that *cis*-interactions of ligands, even to nonconventional receptors, regulate signaling and cancer progression. Furthermore, this *cis*-interaction could be relevant to other contexts, particularly where a reverse gradient of ligands and receptors is observed, such as in stem cell niches. Given that Ephrin-B interacts with EphB for normal stem cell homeostasis, this nonconventional EPHA2/Ephrin-B1 *cis*-interaction, specifically manifested in CSC niches, might serve as an attractive target for therapy, warranting further validation.

## Materials and Methods

### Ethics statement

For the studies involving fresh oral cancer samples, the tumor tissues were collected after getting informed consent from the patients from the Regional Cancer Centre, Thiruvananthapuram. The protocol was approved by the Institute Human Ethical Committee of Rajiv Gandhi Centre for Biotechnology (IHEC/1/2011/04), and Regional Cancer Centre, Thiruvananthapuram (HEC/24/2011). Paraffin embedded tissue blocks for making the tissue microarray (TMA) were collected from the archives of Cochin Cancer Research Centre, Cochin, and the preparation of TMA, IHC staining, scoring and imaging were done at St. Johns Research Institute. The protocol was approved by Institute Human Ethical Committee of all the respective study centers (IEC/150/2023, IHEC/1112022_Exl11). All the animal experiments were conducted adhering to the ARRIVE guidelines, and as per the approved guidelines of the Institute Animal Ethics Committee (IAEC/874/TM/2002, IAEC/875/TM/2002 and IAEC/894/TM/2002) which is under the Committee for the Control and Supervision of Experiments on Animals (CCSEA), Govt. of India. The recombinant DNA works were reviewed and approved by the Institutional Bio-safety Committee (71/IBSC/TTM/2022, 72/IBSC/TTM/2022).

### Antibodies

The antibodies, p-Raf-1, p38 MAPK (MAPK 1/3), TUBB, TUBB2A, TUBA1C, TUBB4B, ERO1L, TUBB3, NDRG1, GAL3, FSCN, GAL1, DSG3, ALDH1A1, Ephrin-B1, EPHA2, EPHB2, mouse IgG and β-Actin were purchased from Santa Cruz Biotechnology, Inc., Dallas, Texas, USA. The other antibodies used were either from Cell Signaling Technology, Danvers, Massachusetts, USA [p-Bad, p-Akt, p-STAT3, pp38 MAPK, SOX-2, OCT4A, EPHA2, Ephrin-B1, pEPHA2 (S897), pEPHA2 (Y772), pEPHA2 (Y594), pEPHA2 (Y588), pEphrin-B (Y324/329), rabbit IgG] or (Thermo Fisher Scientific, Waltham, Massachusetts, USA [pEPHB1/2 (Y594/604), pEphrin-B1 (Y317)]. Peroxidase conjugated AffiniPure Donkey Anti Mouse, AffiniPure Donkey Anti Goat, and AffiniPure Donkey Anti-Rabbit Secondary Antibodies were purchased from Jackson ImmunoResearch Laboratories Inc., West Baltimore Pike, West Grove, PA 19390, USA. Peroxidase conjugated Veriblot for Co-IP experiments was purchased from Abcam, Cambridge, United Kingdom. We used Donkey Anti-Rabbit Alexa Fluor-568, Donkey Anti-Mouse Alexa Fluor-568, Donkey Anti-Rabbit Alexa Fluor-488, Donkey Anti-mouse Alexa Fluor-488, Donkey Anti-Rabbit Alexa Fluor-680, and Donkey Anti-mouse Alexa Fluor-680 (ThermoFisher Scientific, Waltham, Massachusetts, USA) for immunofluorescence and immunohistochemistry experiments.

### Plasmids

pCLXSN-EPHA2 Flag was a gift from the laboratory of Jin Chen (Addgene plasmid # 102755) (23). FUW-ubiquitin-Ephrin-B1-SV40-RFP (biR) was a gift from Eduard Batlle (Addgene plasmid # 65446) (67). pPSmOrange-C1 was a gift from Vladislav Verkhusha (Addgene plasmid # 31899) (37). mT-Sapphire-C1 was a gift from Michael Davidson and Robert Campbell (Addgene plasmid # 54569) (35).

### Cell lines

The cell lines RCB1015, RCB1017, HSC-4, and HSC-3 were obtained from Riken Cell Bank, Japan. All these cell lines were maintained in Dulbecco’s Modified Eagle’s Media with 10% FBS. HSC-4 and HSC-3 cells with stable expression of ALDH1A1-DsRed2 were maintained in 20µg/mL geneticin-containing medium. Cells stably infected with LV-shRNA particles were maintained in 100ng/mL puromycin-containing medium.

### Monolayer and sphere culture

Sphere cells were grown in suspension using Opti-MEM (1X) (Thermo Fisher Scientific, Waltham, Massachusetts, USA), reduced serum medium supplemented with 1X N-2, 20µg/ml EGF and bFGF, and 1X Insulin-Transferrin-Selenium (sphere medium) on a non-adherent dish (coated with 1% Poly (2-hydroxyethyl methacrylate) or polyHEMA to deplete adherence of the dish). All the supplements mentioned above are purchased from Thermo Fisher Scientific, Waltham, Massachusetts, USA. Monolayer culture was maintained in DMEM (Thermo Fisher Scientific, Waltham, Massachusetts, USA) in the presence of 10% FBS (Thermo Fisher Scientific, Waltham, Massachusetts, USA).

### Soft agar colony formation assay

The assay was performed using both monolayer and sphere culture. 5000 cells were seeded for the analysis. After 21 days, the colonies were counted at 10X magnification, and number of colonies/10 fields were plotted.

### RNA isolation, RT-PCR (Reverse transcription PCR) and qRT-PCR

We used the RNeasy mini kit (Qiagen, Hilden, Germany) for RNA isolation from the oral cancer cells. Then, the isolated RNA was quantified using NanoDrop for cDNA preparation. A reaction mix containing RNA (3.5 µg), 5X RT buffer (1X), DNTP (0.1mM), random hexamer primer (10µM), reverse transcriptase enzyme, and RiboLock (reagents for reverse transcription were purchased from ThermoFisher Scientific, Waltham, Massachusetts, USA) was used to reverse transcribe total RNA to cDNA. Further, we performed PCR to check gene expression using gene-specific primers and separated DNA bands using agarose gel for electrophoresis. Then, bands were visualized in Gel Doc EZ imager (Bio-Rad Laboratories, Hercules, California, USA). Similarly, we prepared cDNA and performed qPCR using Power SYBR^TM^ Green PCR Master Mix (ThermoFisher Scientific, Waltham, Massachusetts, USA). Primers for *EPHA2* are F: 5’- TTACCGCAAGAAGGGAGACT-3’, R: 5’-GACAGCGTCTGGAATTCGT-3’. Primers for *EFNB1* F: 5’-GGCAAGATCCCAATGCTGTG-3’, R: 5’-GTTCACAGTCTCATGCTTGCC-3’, Primers for *β-Actin* are F: 5’-CCTTCCTTCCTGGGCATGG-3’, R: 5’- CGCTCAGGAGGAGCAATGA-3’. qPCR results obtained as cycle threshold (Ct) values. The gene expression of specific gene was normalized with expression of *β-Actin* followed with fold change calculation using 2^^-ΔΔCt^ method.

### SILAC labeling

For SILAC labeling, HSC-4 monolayer cells were grown in arginine and lysine-free DMEM (ThermoFisher Scientific, Waltham, Massachusetts, USA) supplemented with heavy arginine and lysine (^13^C_6_ Arg/^13^C_6_Lys;+6Da and +4 Da). SILAC-labeled cells were grown as monolayer cultures while HSC-4 sphere cells were maintained in sphere medium as mentioned above.

### Membrane preparation for LC-MS/MS analysis

The sample preparation and proteomic analysis were done as reported before (68). The cell lysis was done in a high salt buffer (2M NaCl, 10mM HEPES, pH 7.5, 1mM EDTA, pH 8.0) using a polytron tissue homogenizer. The supernatant collected after the centrifugation of the lysate at 250,000 g for 45 min at 4°C was discarded, and the pellet was re-suspended in 0.1 M Na2CO3, pH 11.0, using a polytron homogenizer, and left on ice for 45 min. The high salt lysis and Na2CO3 wash were repeated an additional two times, and the pellet containing the membranous material was used for LC-MS/MS analysis using hybrid LTQ–OrbitrapVelos ETD.

### Cell lysis, Trypsin digestion, phosphopeptide enrichment for LC-MS/MS analysis

The sample preparation was carried out as described before (69). Briefly, the monolayer cells were scraped in PBS and centrifuged to get the cell pellet. Sphere cells were pelleted in the sphere medium and then washed with PBS. Cell pellets were lysed in 9M urea lysis buffer by sonication. The lysates were centrifuged at 12,000 g for 15 min, and the cleared supernatant was collected. The protein concentration of the lysates was estimated using the BCA method, and 3 mg protein from both monolayer and sphere cultures was pooled. Proteins were reduced using 5mM dithiothreitol at 56°C for 45 min. Proteins were further alkylated using iodoacetamide at a final concentration of 10mM and incubated the lysate at room temperature in the dark for 30 min. Lysates were diluted to obtain a final urea concentration of <2 M using HEPES buffer. Sequencing grade trypsin was added to achieve an enzyme to substrate ratio of 1:20 (w/w). Trypsin digestion was carried out overnight at 37°C. The peptide digest was acidified using trifluoroacetic acid (TFA) at a final concentration of 1%. The acidified peptide digest was centrifuged at 4,500 g for 15 min, and the supernatant was desalted using a C18 Sep-Pak cartridge. The desalted peptide digest was frozen and lyophilized for 48 hrs. The lyophilized peptides were reconstituted in 7mM Triethylammonium bicarbonate buffer (pH 9). Peptides were fractionated into 12 fractions using basic reverse phase chromatography. The samples were dried at 4°C and reconstituted in 100 µl of 5% 2, 5-dihydroxybenzoic acid (DHB). TiO_2_ beads (GL Sciences, Inc., Japan Catalogue # 5020-75010) were washed in 5% DHB solution for 15 min. For each fraction, 2 µl of beads was added to achieve the TiO_2_: peptide ratio of 1:1 and incubated on a rotator for 15 min at room temperature. The vials were centrifuged at 1,500 g for 1 min, and the supernatant with unbound peptides was discarded. The beads were reconstituted in 5% DHB solution and loaded on to C8 Stage Tip. The beads were washed twice with 100 µl of 80% acetonitrile with 1% TFA. The phosphopeptides were eluted from TiO_2_ beads using 30 µl of 2% NH_4_OH solution, pH 10.5 (prepared by dissolving 20 µl of NH_4_OH in 1 ml of 40% ACN) into collection tubes containing 20 µl of 0.1% TFA. Then, the phosphopeptides were analyzed using hybrid LTQ–OrbitrapVelos ETD.

### LC-MS/MS analysis

Samples were analyzed on a hybrid LTQ–Orbitrap Velos (Thermo Scientific, Bremen, Germany) mass spectrometer equipped with a nanoelectrospray ion source interfaced with an Easy nano-LC (Thermo Scientific, Bremen, Germany) for sample delivery. The samples were loaded on to a 2 cm trap column (75 µm ID; Magic C 18 AQ, 5µm, 100 Å, Michrom Bioresources Inc.) using 1% formic acid with a flow rate of 3 µl/min and further resolved on a 15 cm analytical column (75 µm ID; Magic C 18 AQ, 3µm, 100 Å, Michrom Bioresources Inc., USA) using a 100-minute gradient from 5% to 30% acetonitrile in 0.1% formic acid with a flow rate of 350 nl/min before introducing into the mass spectrometer. The spray voltage was set to 2.2 kV while the capillary temperature was set to 250°C. The MS instrument was operated in data-dependent acquisition mode. A survey scan from m/z 350–1,700 was acquired in the Orbitrap with resolution 60,000 with a maximum AGC target value of 1,000,000 ions. The fifteen most intense peptide ions with charge states ≥2 were sequentially isolated to a target value of 50,000 ions and fragmented in the higher-energy collisional dissociation (HCD) cell using 39% normalized collision energy. The maximum ion injection times for MS and MS/MS were set to 100 ms and 200 ms, respectively. Fragment ion spectra were detected in the Orbitrap mass analyzer with a resolution of 7,500. For all measurements with the Orbitrap detector, a lock-mass ion from ambient air (m/z 445.120025) was used for internal calibration.

### Data analysis

The mass spectrometry-derived data were searched against the Human RefSeq database (Version 52) using SEQUEST and Mascot search algorithms through Proteome Discoverer software (Version 1.4.1.14) (Thermo Scientific, Bremen, Germany). The protease was specified as trypsin and a maximum of 1 missed cleavage was allowed. Carbamidomethylation of cysteine was set as a fixed modification while oxidation of methionine, phosphorylation at serine, threonine, and tyrosine, and SILAC-labelling (^13^C_6_) at lysine and arginine were set as variable modifications. MS and MS/MS mass tolerances were set to 10 ppm and 0.05 Da, respectively. Peptide identification was based on a 1% false discovery rate calculated using target-decoy database searches. The SILAC ratio for each peptide-spectrum match (PSM) was calculated by the quantitation node and the peptides with ratios greater than two-fold were considered as differentially regulated and were considered for further bioinformatics analysis.

### Bioinformatic analysis

The phosphoproteins identified with <0.5 fold and >2 fold were considered as regulated molecules. The two biological replicates were analyzed separately for pathway enrichment using Genespring Version B13.0. A single Experiment Analysis with 700 source pathways identified the regulated pathways and the molecules regulated in each pathway. All the pathways with p<0.05 were considered as significantly regulated. A Venn diagram of the significantly regulated pathways from the biological replicates identified consistently regulated pathways. The list of regulated molecules in each pathway was used to study the overlap of pathways.

### Western blotting

Cell lysate was prepared using lysis buffer (0.15 M Sodium chloride, 0.025M Tris, 0.001 M EDTA, 0.5% (v/v) NP40, 0.5% (v/v) Triton X 100, 5% glycerol (v/v), 1× protease inhibitor, pH 7.4). The tumor lysate was prepared using RIPA buffer (150mM NaCl, 1% (v/v) NP40, 0.5% (v/v) Sodium deoxycholate, 1% (v/v) SDS, 50mM Sodium orthovanadate, 10mM Sodium Fluoride). The proteins were resolved using SDS-PAGE, followed by the wet-transfer of proteins to a nitrocellulose membrane. After a 10 minutes wash with TBST (500 mM Tris, 1.5M NaCl, 0.15% (v/v) Tween 20), the membrane was blocked using 3% BSA or 10% milk for 1 hour. Then, the membrane was incubated with the primary antibody overnight at 4 °C. The next day, the membrane was washed thrice with TBST to remove unbound primary antibody and incubated in secondary antibody for 1 hour at room temperature. After three times washing with TBST, the membrane was developed using ECL reagent (Bio-Rad Laboratories, Hercules, California, USA).

### Immunofluorescence (IF)/Immunohistochemistry (IHC)

Oral cancer cells were grown as monolayer or spheres for 3 days on confocal dishes. Then, the cells were washed with PBS and fixed using 4% paraformaldehyde (PFA) for 10 minutes. The cells were permeabilized and blocked with 2% fetal bovine serum (FBS) in PBS for surface proteins or with 2% FBS in PBST, which contained 0.3% (v/v) Triton X-100 for intra-cellular molecules for 0.5-1 hour. After removing blocking, cells were incubated with the primary antibody overnight 4 °C. The next day, cells were washed and incubated with the secondary antibody for 1 hour. Then, nuclei were stained using DAPI and washed with PBS to remove excess antibody and DAPI. Further, cells were mounted with ProLong™ Gold Antifade Mountant (ThermoFisher Scientific, Waltham, Massachusetts, USA) and imaged with a confocal microscope, Olympus FV3000 or Nikon A1R LSCM.

Oral cancer patient samples were collected and fixed with 4% PFA for 1 day, followed by storage in 30% sucrose solution. Then, 5µm sections were taken using the LeicaCM1850UV Cryostat. The antigen unmasking was done using 0.01M Citrate buffer, pH 6.0. The IHC staining protocol was similar to IF that used for cultured cells.

### Fluorescence Activated Cell Sorting (FACS)

Cells were trypsinized and resuspended the pellet in DMEM with 10% FBS. Then, the cells were kept inside a CO2 incubator for 1 hour for re-expression, when analyzed for membrane markers. Then, cells were pelleted and fixed with 4% PFA for 10 minutes. After PBS wash, cells were incubated with Alexa fluor-488 tagged primary antibodies for 1 hour at room temperature. Then, after PBS wash, re-suspended the pellet in PBS, filtered with a cell strainer, and used for FACS analysis using FACS AriaIII.

Oral cancer cells stably expressing ALDH1A1-DsRed2 were used for analyzing CSCs, as reported before (32, 33). Briefly, the stable cells were selected using G418 selection. The 2-4% of CSC reporting cells were sorted out, and the enriched population was maintained in DMEM containing 20ng/ml G418. Oral cancer cells stably expressing ALDH1A1 DsRed2 (reporter cell lines) with or without EPHA2/ Ephrin-B1 knock-down were trypsinized and re-suspended the pellet in PBS, filtered with a cell strainer, and analyzed in FACS AriaIII.

### Co-Immunoprecipitation (Co-IP) experiments

For Co-IP, we conjugated protein A/G bead (ThermoFisher Scientific, Waltham, Massachusetts, USA) with the specific antibody or IgG antibodies for 5 hours. We prepared cell lysate using lysis buffer (0.15 M Sodium chloride, 0.025M Tris, 0.001 M EDTA, 0.5% (v/v) NP40, 0.5% (v/v) Triton X 100, 5% glycerol (v/v), 1X protease inhibitor, pH 7.4). Cell lysate was quantified using DC protein assay (Bio-Rad Laboratories, Hercules, California, USA) and 1 mg precleared cell lysate was added to bead-antibody conjugate or bead-IgG conjugate and incubated overnight at 4 °C. The next day, the protein-antibody complex was eluted using 5X loading dye for western blotting experiments or eluted after overnight incubation with IP elution buffer (0.15M Sodium chloride, 0.2M Tris, 0.002 M EDTA, 1% (v/v) Triton X 100, 0.01% (v/v) SDS, 1X protease inhibitor, pH 8.0) for *in vitro* kinase assay.

### ShRNA lentiviral particles and infection

The lentiviral particles for Scrambled (shRNA Control), EPHA2, or Ephrin-B1 shRNA were purchased from Santa Cruz Biotechnology, Inc., Dallas, Texas. These lentiviral particles contain a mixture of three target-specific constructs that encode 19-25 nucleotides (plus hairpin) shRNA designed to knock down the expression of that specific gene. One day before infection, cells were seeded in a 24-well plate. On the day of infection, fresh medium containing 20ug/ml sequabrene was added to cells. Then, 6-10×10^4^ *ifu/ml* of viral shRNA particles were directly added to that medium. This infection procedure was repeated the next day as well. Then, the LV-shRNA-infected cells were selected using puromycin antibiotic (200 ng/ml).

### FRET facilitated photoswitching

PSmOrange-C1 and mT-Sapphire-C1 plasmids were used as pairs for FRET facilitated photoswitching. Then, PSmOrange-C1-EPHA2 or mT-Sapphire-C1-Ephrin-B1 were cloned by amplifying *EPHA2* from pCLXSN-EPHA2 Flag plasmid or *EFNB1* from FUW-ubiquitin-Ephrin-B1-SV40-RFP (biR) plasmid using PrimeSTAR® Max Premix (2X) DNA polymerase mix (Takara Bio Inc., Kusatsu, Shiga, Japan), respectively, using primers with restriction digestion sites, SalI and SmaI. Primers for *EPHA2* were F: 5’-GCATCCCGGGTTAGTCGTCATCCTTG-3’, R: 5’- ATATGTCGACTACCTCGAGATGGAGC-3’. Primers for *EFNB1* were F: 5’ ACATGTCGACCAGGATCTCCACATGGC-3’, R: 5’- ACATCCCGGGCGCGATATCTCAAAC- 3’. Then, the backbone and the PCR products were cut and eluted from the gel using the Gel and PCR clean up kit (Macherey-Nagel, Düren, Germany). Further, they were used for ligation reaction using T4 DNA Ligase (ThermoFisher Scientific, Waltham, Massachusetts, USA) for overnight at 4 °C. Then, the ligated product was transformed into DH5α *E. coli* cells. Then, positive clones were confirmed using restriction digestion and sequencing. Further, HSC-4 cells were transiently transfected with PSmOrange C1-EPHA2 and mT-Sapphire C1-Ephrin-B1 plasmids using Lipofectamine-3000 (ThermoFisher Scientific, Waltham, Massachusetts, USA). Then, these cells were grown in sphere condition for 3 days. Then, we imaged FRET facilitated photoswitching in live cells where we performed a sequential acquisition, as Ex. 639 nm/ Em. 650/25 nm, Ex. 560 nm/ Em. 575/30 nm, Ex. 405 nm/ Em. 488/40 nm. Further, a pulse of 30 iterations of excitation at 405 nm was given. Then, the next cycle was Ex. 639 nm/ Em. 650/25 nm, Ex. 560 nm/ Em. 575/30 nm.

### Proximity Ligation Assay (PLA)

We used Oral cancer cells were grown as spheres for 3 days in confocal dishes. PLA was performed using Duolink® In Situ Red Starter Kit Mouse/Rabbit (Sigma-Aldrich, St. Louis, Missouri, USA). Briefly, the cells were washed with PBS and fixed using 4% paraformaldehyde (PFA) for 10 minutes. Followed by fixation, the cell membrane was stained with wheat germ agglutinin (WGA) Alexa fluor 488 (5μg/ml). Then, cells were permeabilized with 0.5% fetal bovine serum (FBS) in PBST, which contained 0.3% (v/v) Triton X-100 for 30 minutes. Then, cells were washed with PBS and the added the blocking solution. After removing blocking, cells were incubated in primary antibodies (raised in mouse and rabbit, respectively) overnight at 4 °C. The next day, cells were washed and incubated with the PLA probes for 1 hour at 37 °C. Then, PLA probes were removed, and cells were washed with PBS. Subsequently, cells were incubated with ligase for 30 minutes, followed by incubation with polymerase for 1 hour 40 minutes at 37 °C. Further, cells were mounted with mounting medium containing DAPI and imaged with a confocal microscope, Olympus FV3000.

### *In vitro* kinase assay

The Universal Kinase Activity Kit (R&D Systems, Minneapolis, Minnesota, USA) contains ATP, coupling phosphatase (CD39L2), ADP, and malachite green reagents. The assay was performed with mixing varying concentration of EPHA2 kinase obtained from eluate of the co-immunoprecipitation with either Anti-EPHA2, IgG or bead-only control and a small stretch of synthesized peptides with specific tyrosine residues (S BioChem, India) which acted as the acceptor (2mM), ATP (1mM) and CD39L2 (10 ng/μl) followed with 10 minutes’ incubation at room temperature. Further, malachite green reagents were added, which form a green color upon kinase activity, which was read at 655 nm using an ELISA Plate Reader (Bio-Rad Laboratories, Hercules, California, USA). Negative control for the kinase assay lacked the kinase (IP eluate). Peptides used were YVDPHTYEDPNQ, which contains autophosphorylation residues (Y588 and Y594) of EPHA2, GDYGHPVYIVQEMPPQ, which contains Y324 and Y329 residues of Ephrin-B1, and NYCPHYEKVSG, which contains Y317 residue of Ephrin-B1.

### Extreme Limiting Dilution Assay (ELDA)

For *in vitro* sphere formation assay, cells were trypsinized and the serial dilutions were prepared and seeded in non-adherent 96-well plate (coated with 1% Poly (2-hydroxyethyl methacrylate) or polyHEMA to deplete adherence of the plate) in sphere medium and maintained for 6 days, supplementing medium every other day. On the 6^th^ day, counted the spheres in each well and data was used for Extreme limiting dilution assay (ELDA, (https://bioinf.wehi.edu.au/software/elda/). CSC frequency was calculated using ELDA analysis, which is a web-tool available online, and designed by the Bioinformatics Division, The Walter and Eliza Hall Institute of Medical Research in Australia. ELDA plotted a log-fraction plot, the slope of which gives CSC frequency.

NOD.CB17-Prkdc^scid^I12rg^tm1Wjl^/SzJ (NSG) mice were used for generating tumors for *in vivo* serial dilution xenograft assay, following approved protocols from IAEC. Cells were serially diluted and mixed with Corning® Matrigel® Growth Factor Reduced (GFR) Basement Membrane Matrix and injected subcutaneously on the flanks of the mice. After palpable tumors were formed, the tumors were collected and representative images were taken. CSC frequency was calculated using ELDA.

### Site-directed mutagenesis

Site-directed mutagenesis of tyrosine to phenylalanine in Ephrin-B1 RFP plasmid (Ephrin-B1 and RFP are activated under two different promoters, Ephrin-B1 under UBQ promoter and RFP under SV40 promoter respectively) was done using PfuUltra II Fusion HS DNA Polymerase (Agilent Technologies, Inc., Santa Clara, California, USA) with completely complimentary primers of length between 26-35 base pairs which occupies mutated base at the centre of the primers. Primers for Ephrin-B1 Y323F are F: 5’-GTGAGTGGGGAC TTCGGGCATCCTGTC-3’, R: 5’- GACAGGATGCCCGAAGTCCCCACTCAC-3’. Primers for Ephrin-B1 Y316F are F: 5’- TACTGCCCCCACTTTGAGAAGGTGAGTG-3’, R: 5’- CACTCACCTTCTCAAAGTGGGGGCAGTA-3’. After PCR amplification with the mutated primers, the PCR product was digested with the DpnI enzyme and performed transformation into DH5α E. Coli cells. Positive clones were confirmed with restriction digestion and sequencing reaction.

### Orthotopic models for oral carcinoma

NOD.CB17-*Prkdc^scid^/*J (SCID) or NOD.CB17-Prkdc^scid^I12rg^tm1Wjl^/SzJ (NSG) mice were used for generating tumors following approved protocols from IAEC. For the *in vivo* serial dilution xenograft assay, 1x10^6^ cells of oral cancer cells were mixed with Corning® Matrigel® Growth Factor Reduced (GFR) Basement Membrane Matrix and injected subcutaneously on the flanks of the mice. Tumors were collected after 2-3 weeks and photographed. For the orthotopic model, 1x10^6^ cells of oral cancer cells were mixed with Corning® Matrigel® Growth Factor Reduced (GFR) Basement Membrane Matrix and injected into the floor of the mouth region superficial to the mylohyoid muscle. The animals were periodically monitored for overall survival analysis. The weight of the animals and tumor volume were measured periodically (using digital Vernier calipers) for 10 weeks. The death of the animal or ethical endpoint (visible signs of pain or distress, or if the weight loss was >25% of pre-injection) was considered as the survival time.

### *In-silico* analysis of oral cancer patient data

To analyze the pattern of expression of EPHA2, Ephrin-B1, and EPHB2 mRNA in oral cancer specimens, data was downloaded from the Gene Expression Omnibus data portal (https://www.ncbi.nlm.nih.gov/geo/). The GEO accession numbers of the datasets are GSE13601, GSE65858. The platform used for expression profiling in all three datasets were microarray. In GSE65858, a total of 300 samples were considered for analysis and quality control procedures were applied to microarray probe-level intensity files. A total of 255 tumor arrays remained after removing low-quality arrays, duplicate arrays, and arrays from non-HNSCC samples. Expression values were log2-transformed and normalized using RSN. The relative normalized log-transformed microarray data was used for all the analysis. The TCGA data (N=515) was accessed from the TCGA Research Network: https://www.cancer.gov/tcga (accessed on 15 November 2021).

Descriptive statistics were used for all clinical variables. The difference in gene expression levels was evaluated by the Mann–Whitney U test/Kruskal–Wallis test. Correlations were evaluated by Pearson’s rank test. Heatmap analysis was also performed to analyze the association of various genes, EPHA2, Ephrin-B1 and EPHB2. Kaplan-Meier analysis was used to examine the estimated differences in overall survival between the various groups. Log-rank test (Mantel-Cox) was used to compare the survival between groups. Survival analysis was performed by stratifying tumors based on the upper or third quartile expression of EPHA2 and Ephrin-B1 mRNA levels and grouped into tumors with high and low expression. For all tests, a p-value of <0.05 was considered to be statistically significant. All statistical analysis was carried out using the software XLSTAT - version 2022.2.1.

### Tissue microarray (TMA) and IHC analysis

Archival samples were collected from Cochin Cancer Research Centre (CCRC). Patients were categorized as recurrent and disease-free groups where patients with disease-free survival of less than 15 months were considered as recurrent (25 samples) while patients with disease-free survival for more than 15 months were considered as disease-free group (27 samples).

Immunohistochemistry for the antibodies was done on each of the TMA sections as per standard protocol using the Ventana Benchmark^XT^ staining system (Ventana Medical Systems, Tucson, AZ, USA). Briefly, 5 μm thick sections were fixed in a hot air oven at 60°C for 60 min and loaded onto the IHC staining machine. De-paraffinization was performed using EZ Prep solution (Proprietary-Ventana reagent), and antigen retrieval was done using Cell Conditioning solution 1 (CC1) for 60 min. The primary antibody was added manually and incubated for 32 min at room temperature. Optiview DAB Detection Kit (Ventana Medical Systems) was used to visualize the signal, using DAB (3–3′diaminobenzidine) as the chromogen. Further, the sections were automatically counterstained with hematoxylin II (Ventana Medical Systems) for 12 min. The slides were removed from the autostainer, washed in de-ionized water, dehydrated in graded ethanol, cleared in xylene, and examined by microscopy. Appropriate positive and negative controls were run for each batch. Two pathologists scored the staining independently. Nuclear/cytoplasmic staining in more >1% of tumor cells was considered positive. Data from 24 recurrent and 26 disease-free samples were used for the statistical analysis.

## Acknowledgements

We acknowledge Akhilesh Pandey and the Proteomic facility of Institute of Bioinformatics, Bangalore, for the proteomic analyses. The contribution of Annie Agnes Suganya S for Fig 1C is acknowledged. We thank the contributions of the Animal Research Facility, the Confocal Facility and FACS Facility of RGCB. Reshma Raj R received funding from Council of Scientific & Industrial Research, India (09/716(0188)/2019-EMR-I). The work was financially supported by the funding from DST-SERB (SR/S0/HS/0133/2010), DBT (BT/PR14379/Med/30/536/2010), and RGCB LRF.

## Supporting information

**S1 Fig. Sphere culture enriches self-renewing population.**

(A) Serial passaging sphere formation assay with three oral cancer cells as indicated. 3D Spheres were formed up to three passages. The image indicates primary and tertiary spheres formed in the three oral cancer cells. (B) We performed a soft agar colony formation assay with three oral cancer cells grown as monolayer (ML) or 3D sphere (Sph). After 21 days, the colonies were counted at 10X magnification, and the number of colonies/10 fields in monolayer and sphere from 3 independent experiments were plotted as mean ± SD. (C) HSC-4 cells were grown as ML or Sph for 3 days. Then, we stained the cells with ALDH1A1 and Hoechst. (D) HSC-4 cells grown as monolayer or 3D spheres for 3 days were used for RNA isolation followed with RT-PCR (E) 3- Day-old HSC-4 ML or 3D Sph were used for cell lysis preparation followed by western blotting to check expression of OCT4A. Statistical analysis was done by an unpaired-two-tailed students t-test. ***, **** represents p<0.001 and p<0.0001 respectively.

**S2 Fig. Diagrammatic representation of proteomic analyses.**

(A) HSC-4 cells were grown as monolayer in SILAC medium containing heavy arginine and lysine, while HSC-4 3D sphere cells were maintained in sphere medium with normal amino acids for 6 days. The membrane prep was used for LC-MS/MS analysis, as shown in the diagram. (B) The cells were grown as described, and the lysates were used for phosphoproteomic analysis

**S3 Fig. Validation of proteomic data.**

(A) HSC-4 cells were grown in sphere conditions for 3 days, fixed, and stained for the indicated molecules. The scale bar represents 10µm. (B) Some of the molecules obtained from phosphoproteomic analysis were confirmed using western blotting using 6-Day-old monolayer (ML) or 3D Sphere (Sph). (C) The differentially expressed phospho enriched peptides obtained from the phosphoproteomic analysis were analyzed for pathway enrichment. The bar plot represents differential phosphorylation of the molecules in the Eph/Ephrin signaling pathway. (D) HSC-4 Cells were grown as monolayer or sphere for 3 days, and were used for FACS analysis to check surface expression of EPHA2, Ephrin-B1, and EPHB2 under non-permeabilizing condition. FITC-A represents the positivity of the indicated molecules.

**S4 Fig. Western blots of EPHA2 and Ephrin-B1.**

(A) Western blot experiment to show bands of EPHA2 (full blot) of HSC-4 oral cancer cells with knock-down of EPHA2. β-Actin was used as the loading control. (B) Bar plot represents fold change of knock-down over control in western blot experiment. (C) Bar plot represents fold change of knock-down over control in qRT-PCR experiment. (D) Western blot experiment to show bands of Ephrin-B1 in a full blot of HSC-4 oral cancer cells with knock-down of Ephrin-B1. (E) The Bar plot represents the fold change of knock-down over control in the western blot experiment. (F) The Bar plot represents the fold change of knock-down in the qRT-PCR experiment. Statistical analysis of 3 independent experiments was done by a paired-two-tailed students t-test. **, ***, **** represents p<0.01, p<0.001 and p<0.0001 respectively.

**S5 Fig. Western blot of pEphrin-B1.**

(A-B) Western blot experiment to show bands of pEphrin-B (Y324/329) and pEphrin-B1 Y317 (full blot) of 3-Day-old HSC-4 3D spheres with knock-down of EPHA2. β-Actin was used as the loading control.

**S6 Fig. Cloning of Ephrin-B1 Y324F or Ephrin-B1 Y317F.**

(A-B) We performed site-directed mutagenesis of FUW-ubiquitin-Ephrin-B1-SV40-RFP plasmid. The Ephrin-B1 Y324F clone was confirmed by restriction digestion and sequencing, while the parent plasmid was used as the positive control.

**S7 Fig. *In vitro* sphere formation assay (ELDA).**

(A) Western blot confirmation of single knock-down of EPHA2 or Ephrin B1 and dual knock-down of both molecules. (B) We normalized the expression with β-Actin. Then the fold change was calculated by dividing relative expression in knock-down by control, and was plotted as mean ± SD values from 3 biological replicates. (C) The image shows the 6-Day spheres formed in the dilution of 3000 HSC-4 cells. The scale bar represented 100µm. The bar plot shows mean ± SD values of the total number of spheres formed per well for each dilution in single knock-down of EPHA2 in HSC-4 cells. The graph represents a log fraction plot obtained from ELDA analysis, and the slope of it represents cancer stem cell frequency. The table shows the data for ELDA analysis. (D) The bar plot shows mean ± SD values of the total number of spheres formed per well for each dilution in single knock-down and dual knock-down cells of RCB1015. (E) the table shows the data for the ELDA analysis of RCB1015. Statistical analysis other than ELDA was done by a paired-two-tailed students t-test. *, **, **** represents p<0.05, p<0.01 and p<0.0001 respectively.

**S8 Fig. ELDA for serial dilution xenograft and mutants.**

(A-D) Table shows the data for ELDA analysis for the indicated combinations.

**S9 Fig. Cloning of PSmOrange-C1 EPHA2 and mT-Sapphire-C1 Ephrin-B1.**

(A) PSmOrange-C1-EPHA2 clone was confirmed using restriction digestion and sequencing. Restriction digestion was performed with enzymes SalI and SmaI. The expected band sizes were 4697 for the backbone and 2972 for the insert. (B) mT-Sapphire-C1-Ephrin-B1 clone was confirmed using restriction digestion and sequencing. Restriction digestion was performed with enzymes SalI and SmaI. The expected band sizes were 4712 for the backbone and 1067 for the insert.

**S10 Fig. FRET Facilitated photoswitching.**

(A-C) HSC-4 cells were transiently transfected with empty vectors (PSmOrange-C1 and mT-Sapphire-C1) or with the PSmOrange C1-EPHA2 and mT-Sapphire-C1 Ephrin-B1. Then, these cells were grown as spheres for 3 days, followed by imaging for FRET facilitated photoswitching in live cells, where we performed a sequential acquisition, as Ex. 639 nm/ Em. 650/25 nm, Ex. 560 nm/ Em. 575/30 nm, Ex. 405 nm/ Em. 488/40 nm. Further, a pulse of 30 iterations of excitation of 405 nm was given. Then, the next cycle was Ex. 639 nm/ Em. 650/25 nm, Ex. 560 nm/ Em. 575/30 nm. PSmOrange-C1-EPHA2 was given a 488 nm iteration instaed of 405 in one experiment as a positive control for photoswitching. (B) FRET Facilitated photoswitching images of PSmOrange-C1 and mT-Sapphire-C1 empty vectors, which served as a negative control. (C) FRET Facilitated photoswitching images of PSmOrange-C1-EPHA2 and mT-Sapphire-C1-Ephrin-B1.

**S11 Fig. ELDA for *cis-*interaction.**

Table shows data for ELDA analysis of 1:1 mixture of single knock-down of EPHA2 and Ephrin-B1.

**S12 Fig. Western blotting and IHC of xenografts.**

(A) 1.2×10^6^ RCB1015 wild-type or Ephrin-B1 knocked-down cells were injected subcutaneously on the flanks of the SCID mice. Tumors were collected after 28 days, and tumor tissue lysate was prepared and probed for the indicated molecules. (B) 1.2×10^6^ RCB1017 wild-type or EPHA2 knocked-down cells were injected subcutaneously on the flanks of the SCID mice. Tumors were collected after 34 days, and western blot was performed as described. (C) RCB1015 xenograft sections were stained for EPHA2, Ephrin-B, and ALDH1A1. (D-F) Bar plots represent mean intensity per field of EPHA2, Ephrin-B1 or ALDH1A1 in control and knock-down sections. Statistical analysis was done by an unpaired-two-tailed students t-test. *, ***, *** represents p<0.05, p<0.001 and p<0.0001, respectively.

**S13 Fig. Clinical data analyses.**

(A) Univariate analysis depicting distribution of the Ephrin-B1 (*EFNB1*), *EPHA2*, and *EPHB2* mRNA levels in OSCC specimens (N=31) compared to normal samples (N=26), data was downloaded from the Gene Expression Omnibus data portal. The GEO accession number of the dataset was GSE13601. (B) Hierarchical clustering heat map analysis with TCGA mRNA data set of HNSCC (N=515) to check expression of *EFNB1*, *EPHA2*, and *EPHB2*. Each vertical line in the map represents a single patient. (C) Correlation between EPHA2 and Ephrin B1 was checked in TCGA as well as GSE data set using Pearson’s correlation analysis.

**S1 Table. Differential expression of membrane proteins in sphere and monolayer.**

**S2 Table. Differential expression of phosphopeptides in sphere and monolayer in replicate 1.**

**S3 Table. Differential expression of phosphopeptides in sphere and monolayer in replicate 2.**

**S4 Table. Differential expression of phosphoproteins in sphere and monolayer of replicate 1 and 2.**

**S5 Table. Differentially regulated pathways in spheres compared to monolayer from replicate 1 and 2.**

**S6 Table. Differentially regulated pathways implicated in self-renewal.**

**S7 Table. Differentially regulated phosphoproteins of Eph/Ephrin signaling pathway.**

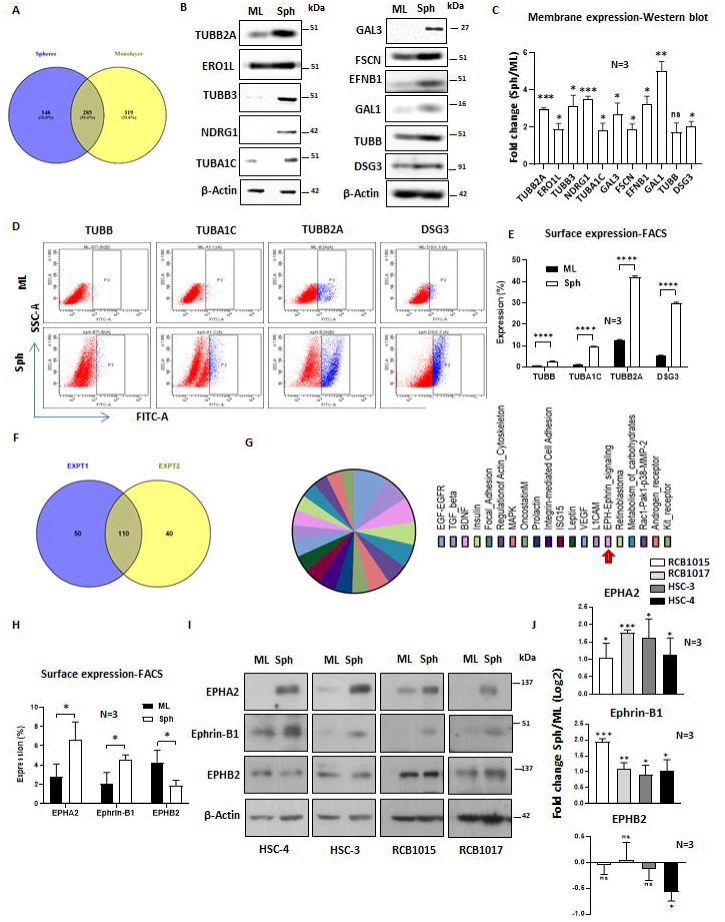

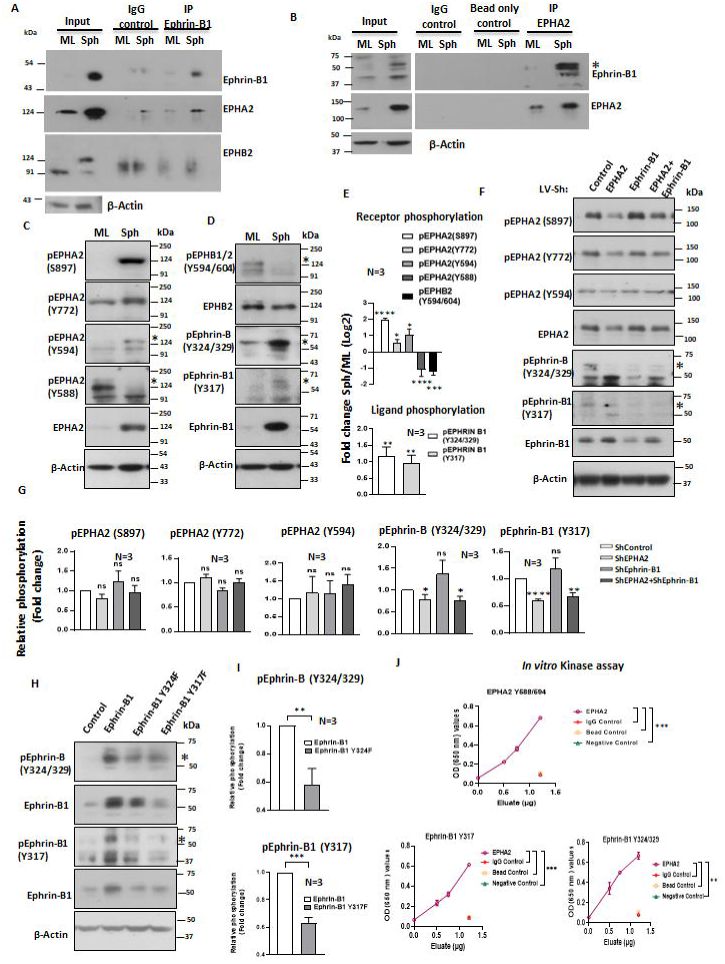

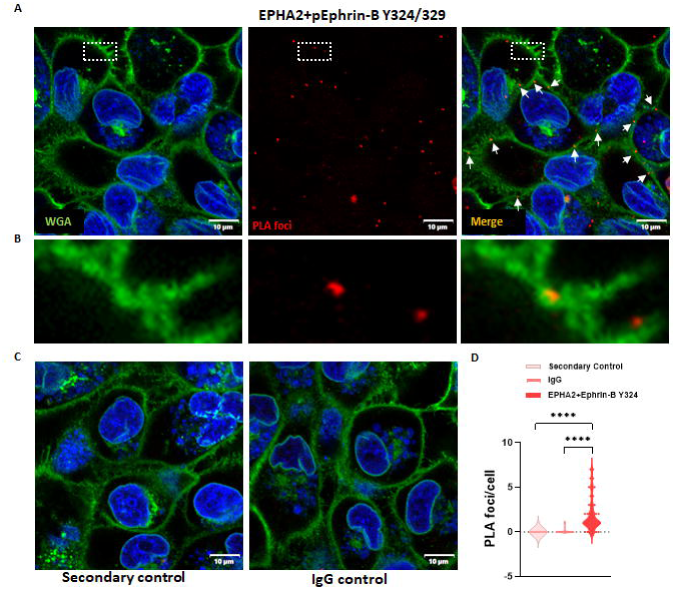

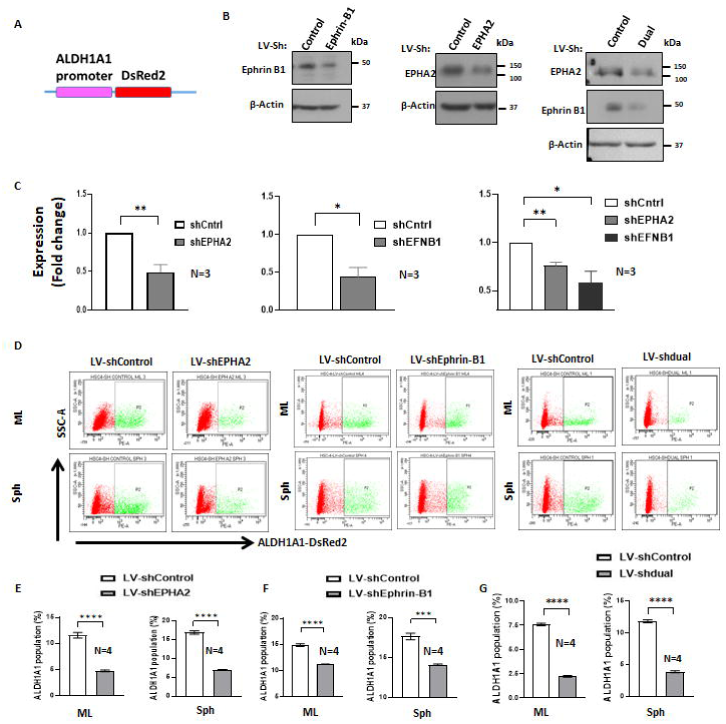

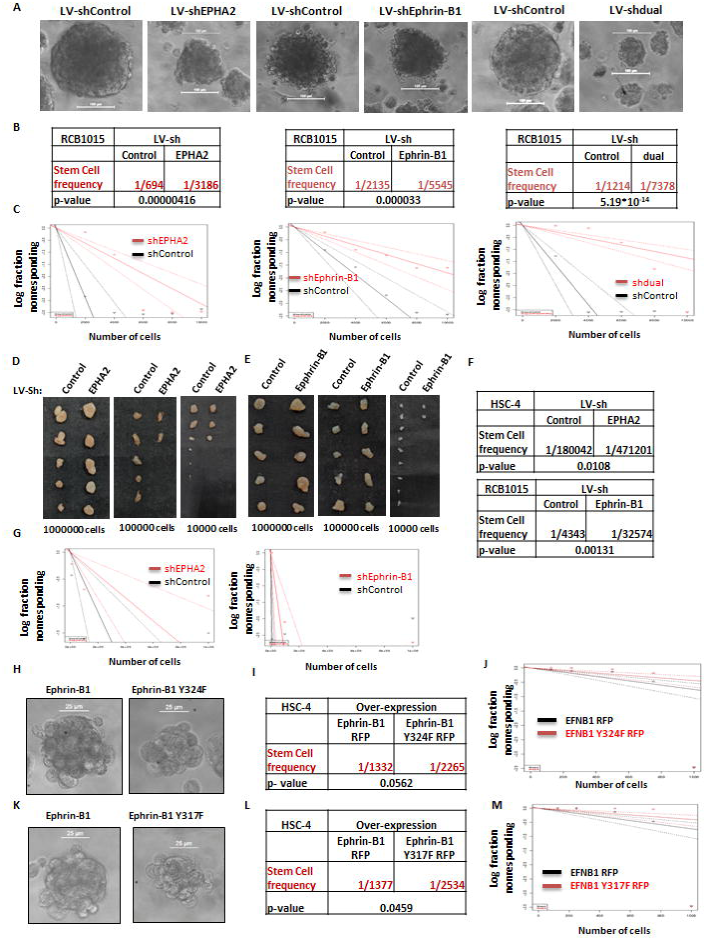

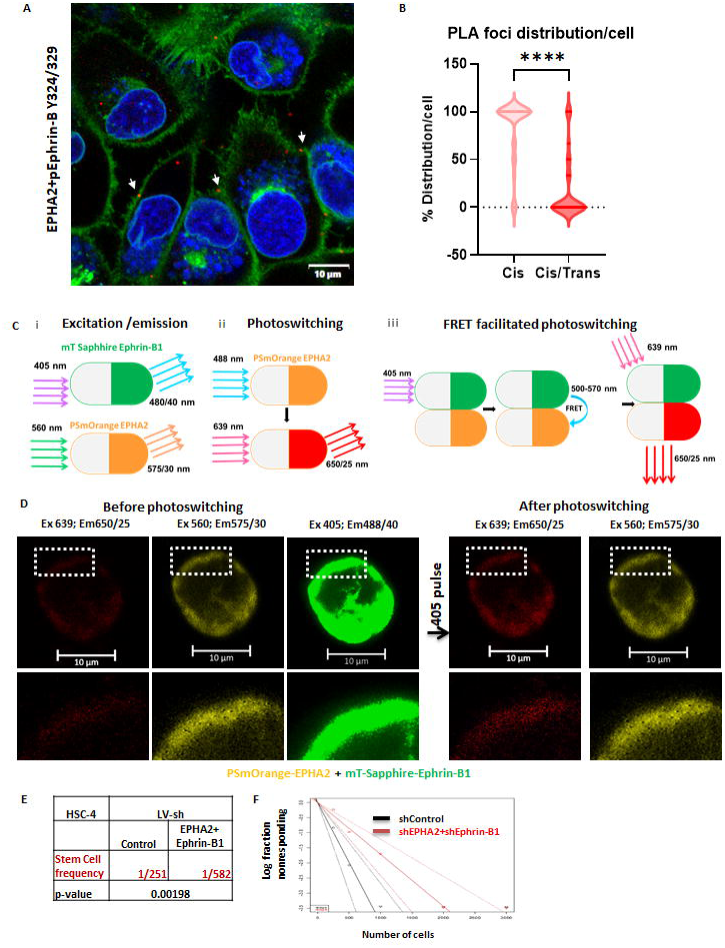

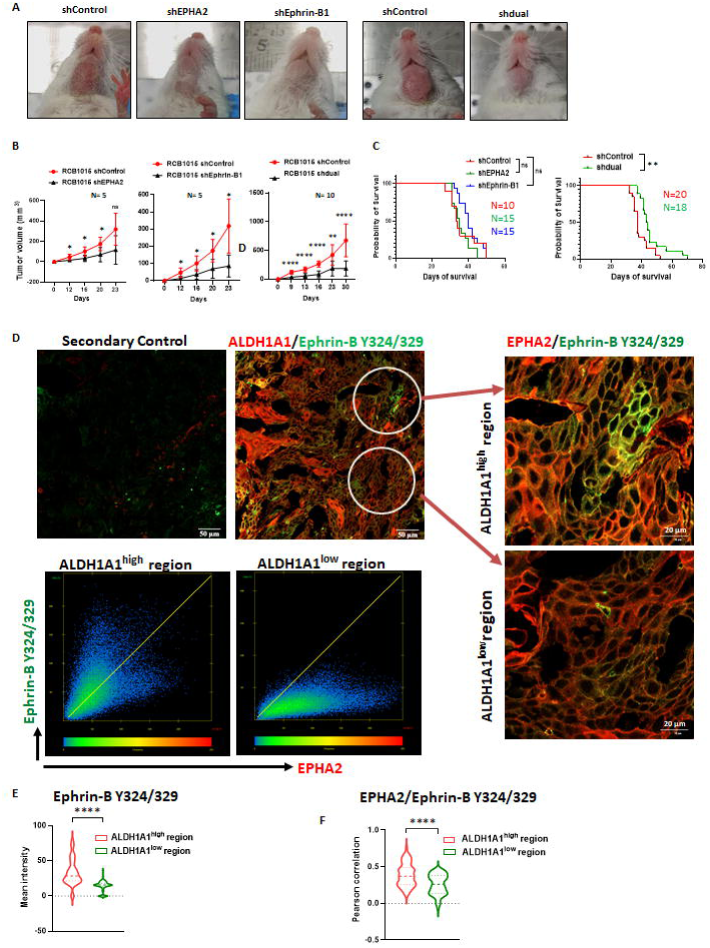

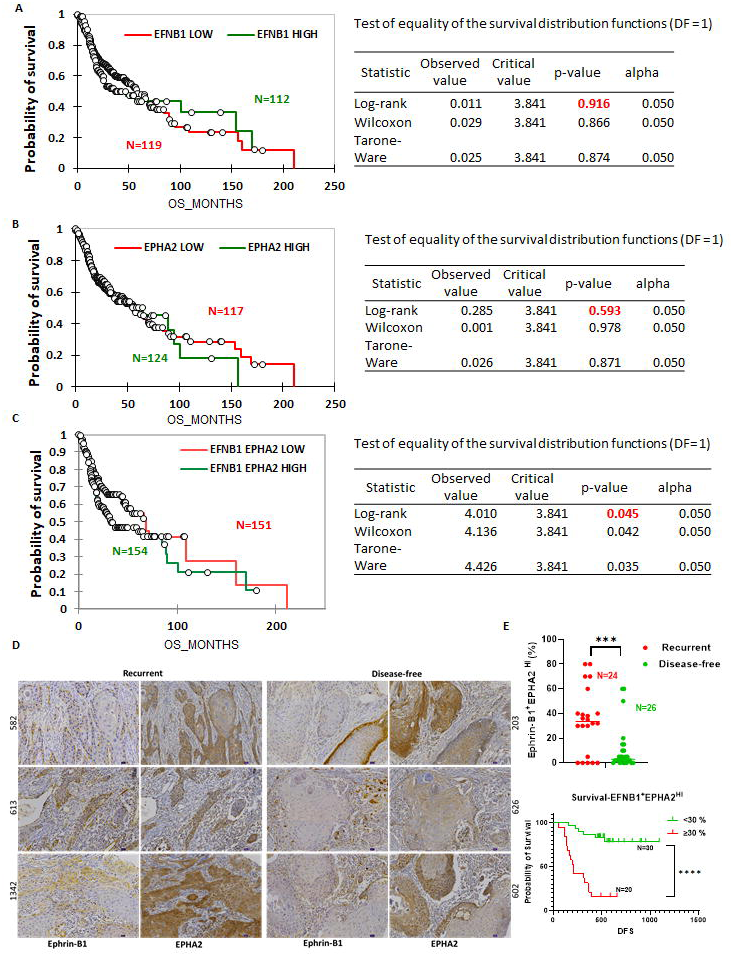

