## Supplementary Figures for "EPHA2-Ephrin-B1 *cis*-interaction supports self-renewal ability leading to recurrence of oral cancer"

### Supplementary Figure 1

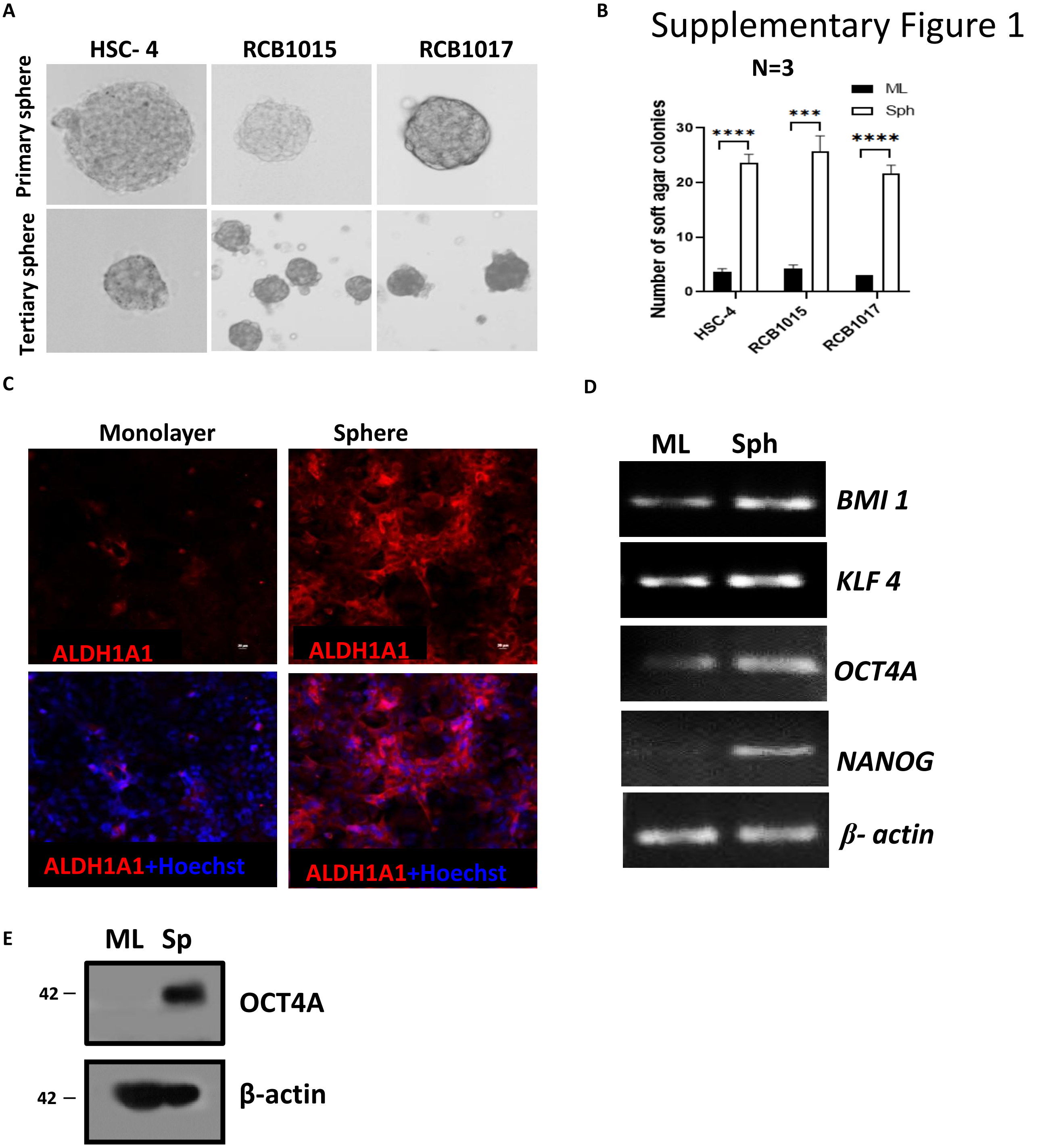

Supplementary Figure 2

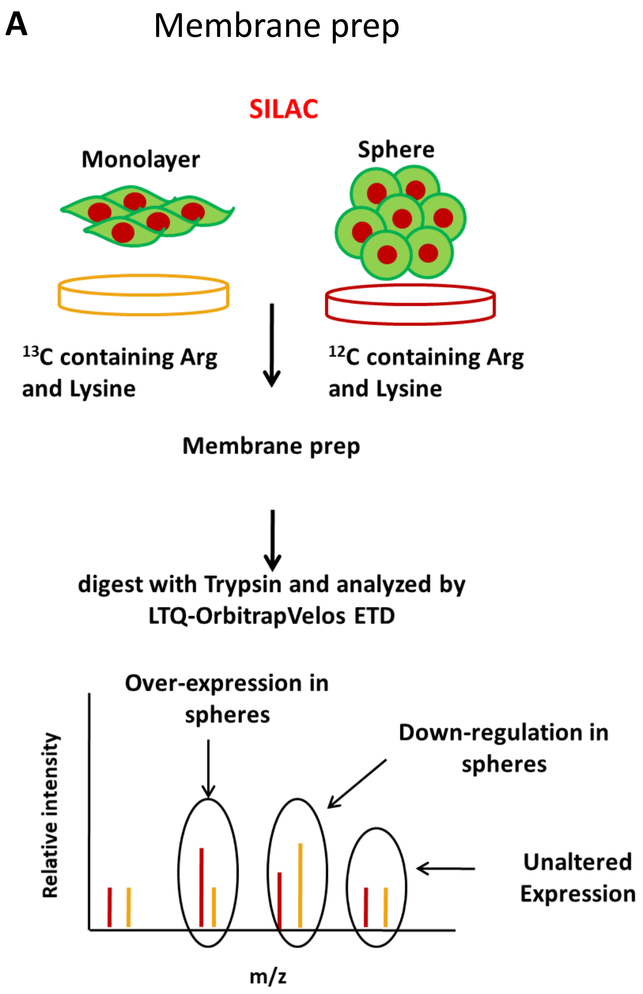


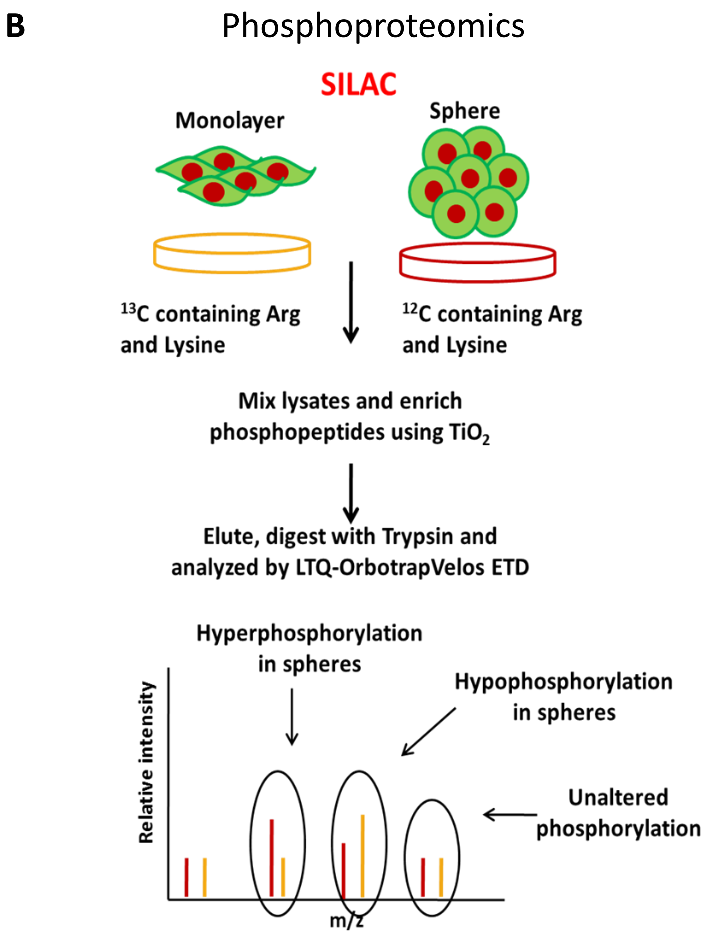

### Supplementary Figure 3

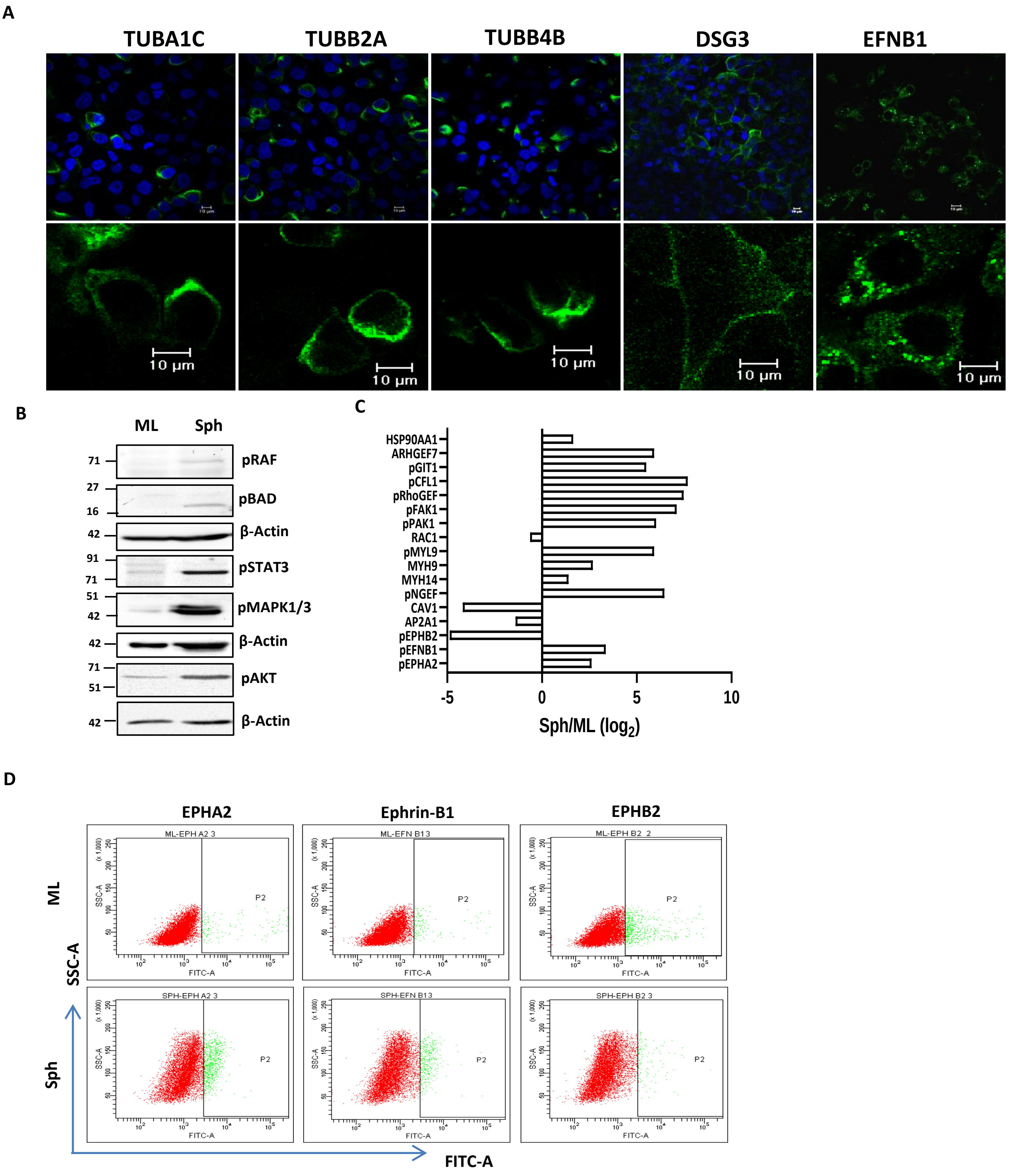

### Supplementary Figure 4

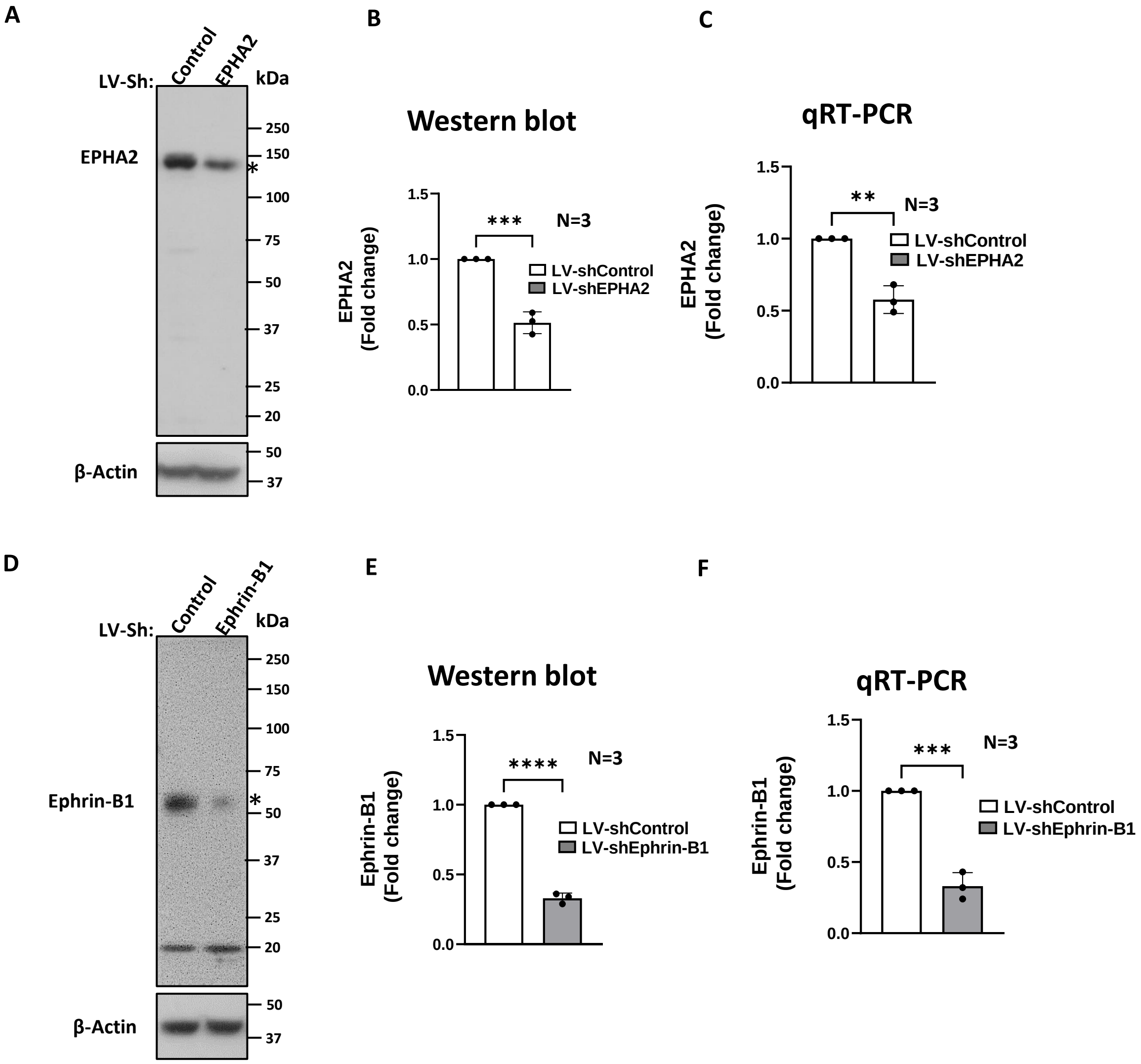

### Supplementary Figure 5

A

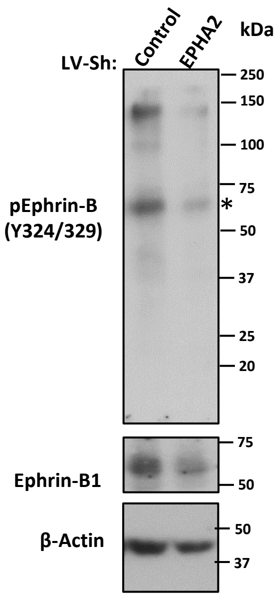


B

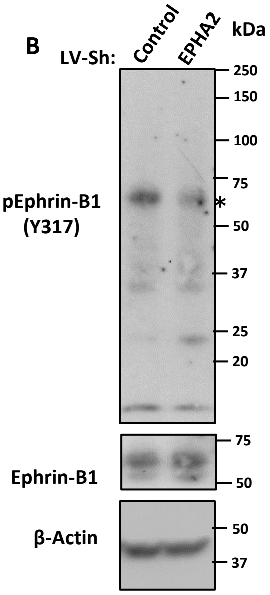

### Supplementary Figure 6

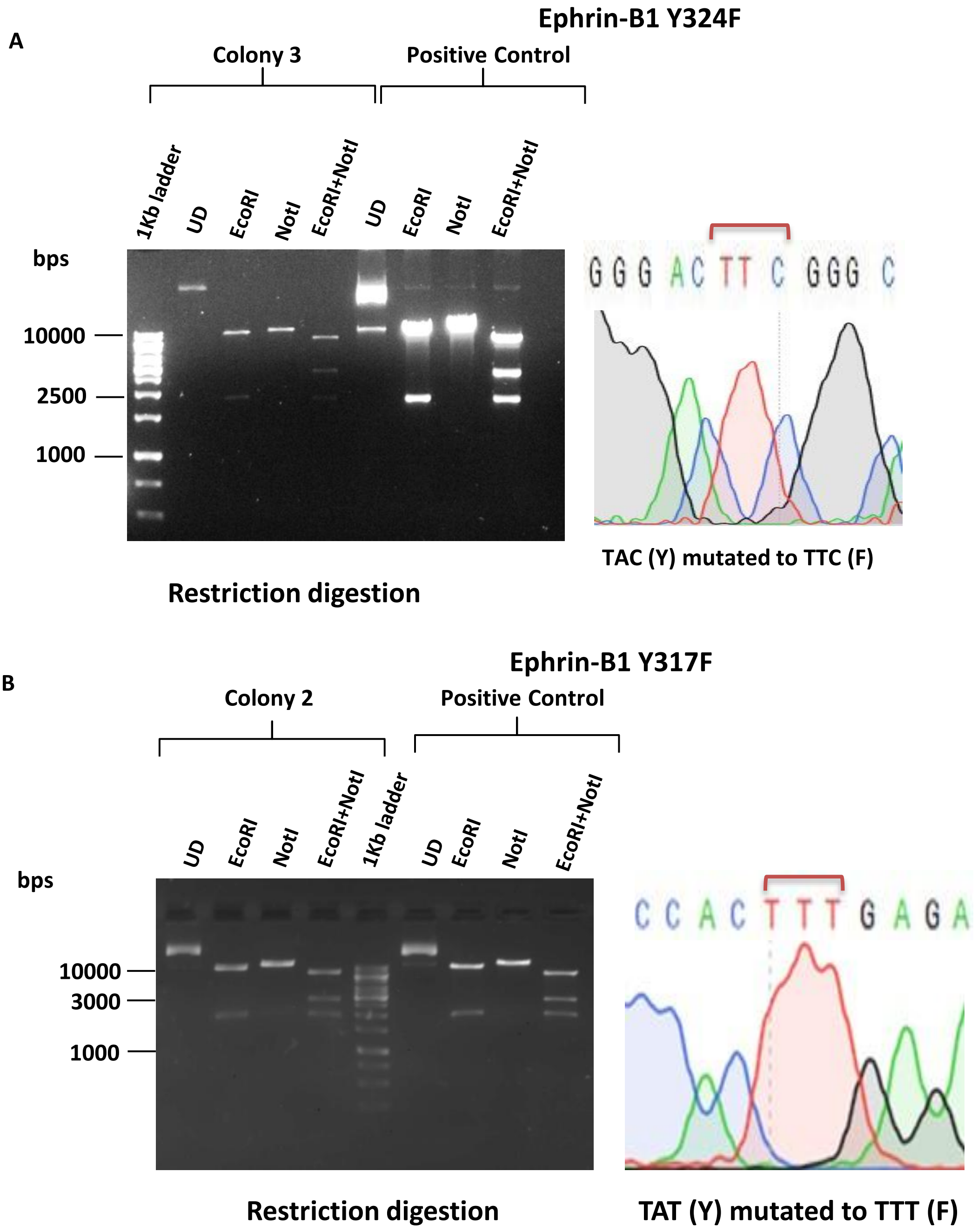

### Supplementary Figure 7

A

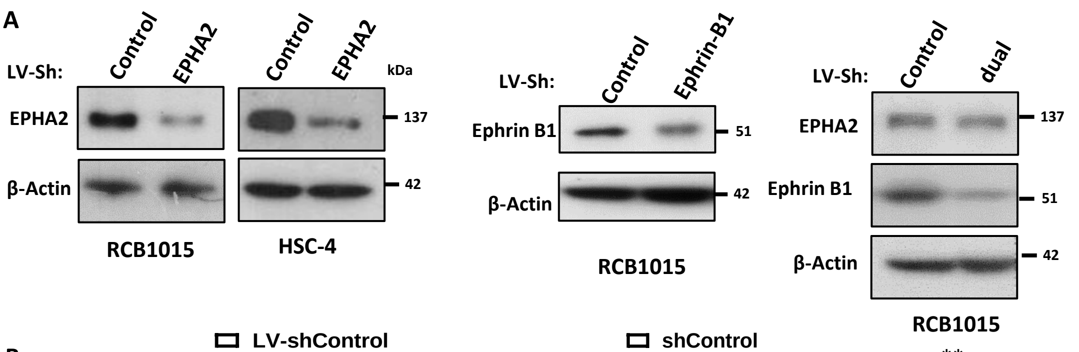


B

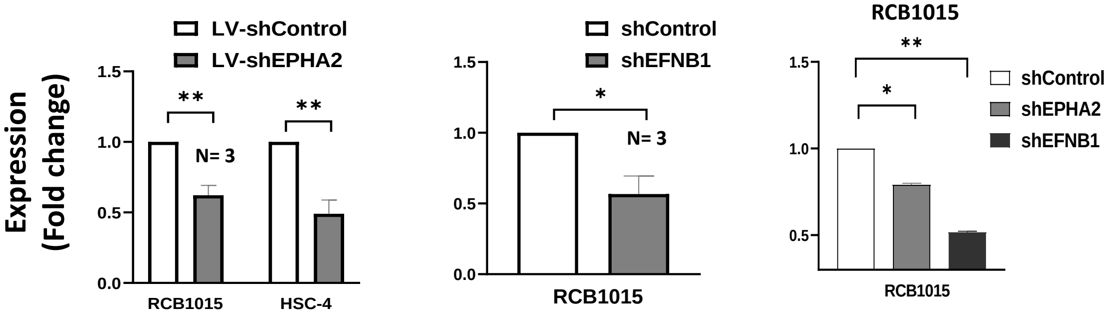


C

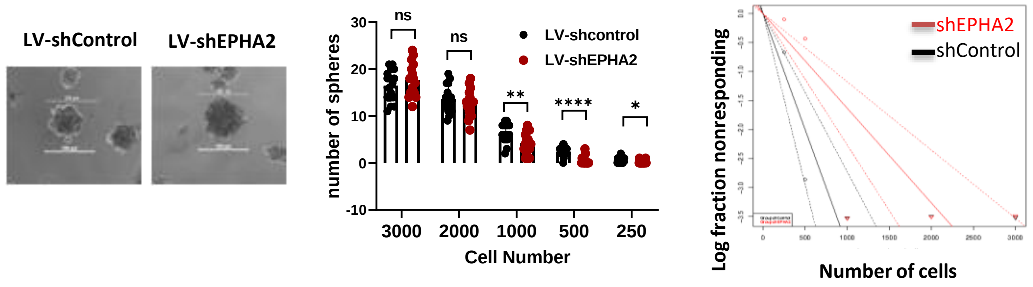


| HSC-4 | LV-sh | |
|---|---|---|
| | Control | EPHA2 |
| Cell No. | | |
| 3000 | 17/17 | 17/17 |
| 2000 | 17/17 | 17/17 |
| 1000 | 17/17 | 17/17 |
| 500 | 16/17 | 6/17 |
| 250 | 8/17 | 2/17 |
| Stem Cell frequency | 1/250 | 1/613 |
| P value | 0.000827 | |

D

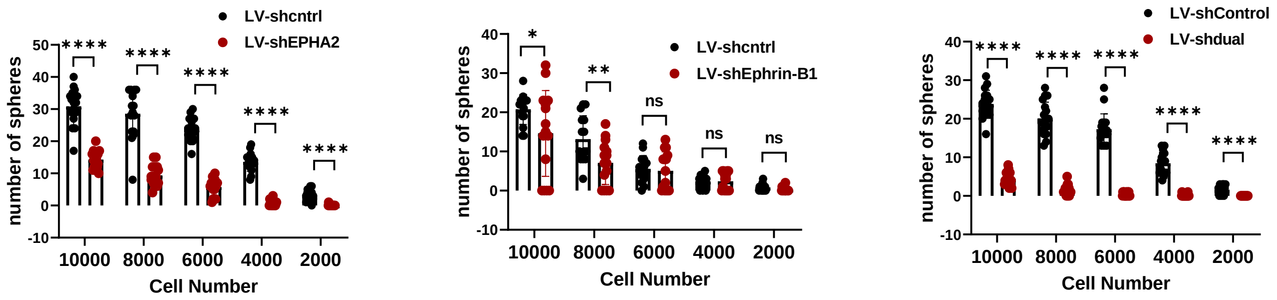


E

| RCB1015 | LV-sh | |
|---|---|---|
| | Control | EPHA2 |
| Cell No. | | |
| 10000 | 16/16 | 16/16 |
| 8000 | 16/16 | 16/16 |
| 6000 | 16/16 | 16/16 |
| 4000 | 16/16 | 10/16 |
| 2000 | 15/16 | 1/16 |
| Stem Cell frequency | 1/694 | 1/3186 |
| P value | 0.00000416 | |

| RCB1015 | LV-sh | |
|---|---|---|
| | Control | Ephrin-B1 |
| Cell No. | | |
| 10000 | 15/15 | 12/15 |
| 8000 | 15/15 | 12/15 |
| 6000 | 14/15 | 10/15 |
| 4000 | 12/15 | 9/15 |
| 2000 | 9/15 | 3/15 |
| Stem Cell frequency | 1/2135 | 1/5545 |
| P value | 0.000033 | |

| RCB1015 | LV-sh | |
|---|---|---|
| | Control | dual |
| Cell No. | | |
| 10000 | 18/18 | 18/18 |
| 8000 | 18/18 | 15/18 |
| 6000 | 18/18 | 9/18 |
| 4000 | 18/18 | 2/18 |
| 2000 | 13/18 | 0/18 |
| Stem Cell frequency | 1/1214 | 1/7378 |
| P value | 5.19*10-14 | |

### Supplementary Figure 8

**A**

| HSC-4 | LV-sh | |
|---|---|---|
| | Control | EPHA2 |
| Cell No. | | |
| 1000000 | 7/9 | 5/9 |
| 100000 | 8/10 | 5/10 |
| 10000 | 7/20 | 4/20 |
| Stem Cell frequency | 1/180042 | 1/471201 |
| P value | 0.0108 | |

**B**

| RCB1015 | LV-sh | |
|---|---|---|
| | Control | Ephrin-B1 |
| Cell No. | | |
| 1000000 | 5/5 | 5/5 |
| 100000 | 5/5 | 5/5 |
| 10000 | 9/10 | 2/10 |
| Stem Cell frequency | 1/4343 | 1/32574 |
| P value | 0.00131 | |

**C**

| HSC-4 | Overexpression | |
|---|---|---|
| | EFNB1 RFP | EFNB1 Y324F RFP |
| Cell No. | | |
| 1000 | 16/16 | 16/16 |
| 750 | 9/24 | 4/24 |
| 500 | 3/24 | 1/24 |
| 250 | 4/40 | 0/40 |
| 125 | 1/16 | 0/16 |
| Stem Cell frequency | 1/1332 | 1/2265 |
| P value | 0.0562 | |

**D**

| HSC-4 | Overexpression | |
|---|---|---|
| | EFNB1 RFP | EFNB1 Y317F RFP |
| Cell No. | | |
| 1000 | 16/16 | 16/16 |
| 750 | 9/24 | 1/24 |
| 500 | 3/24 | 0/24 |
| 250 | 0/16 | 0/16 |
| 125 | 0/16 | 0/16 |
| Stem Cell frequency | 1/1377 | 1/2534 |
| P value | 0.0459 | |

### Supplementary Figure 9

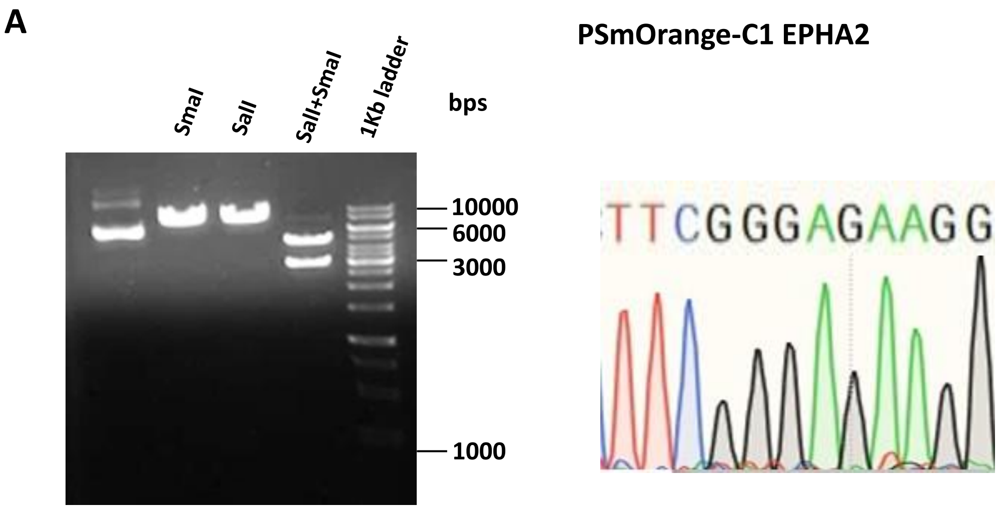


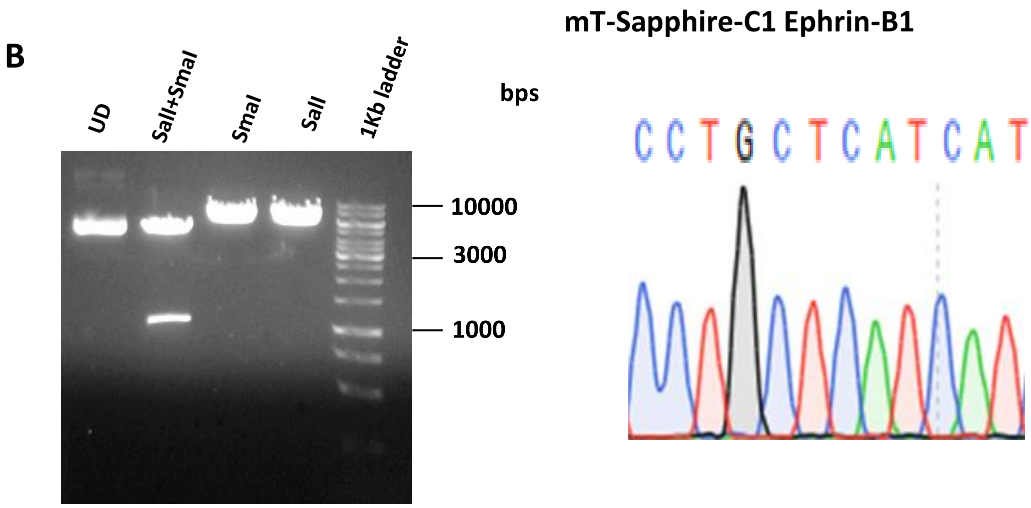

### Supplementary Figure 10

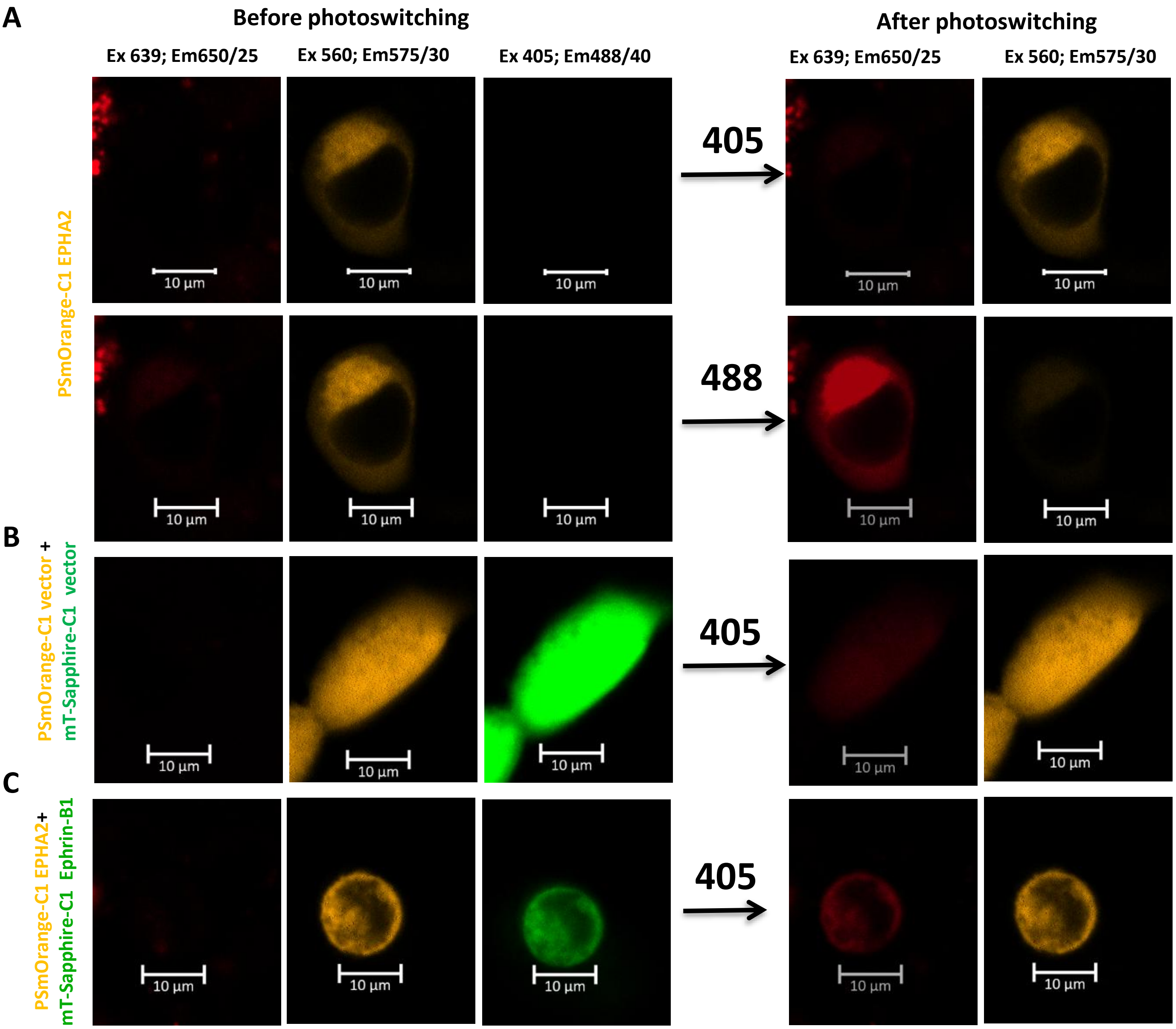

### Supplementary Figure 11

| HSC-4 | LV-sh | |
|---|---|---|
| | Control | EPHA2+ Ephrin-B1 |
| Cell No. | | |
| 3000 | 16/16 | 16/16 |
| 2000 | 16/16 | 16/16 |
| 1000 | 16/16 | 13/16 |
| 500 | 14/16 | 10/16 |
| 250 | 9/16 | 3/16 |
| Stem Cell frequency | 1/251 | 1/582 |
| P value | 0.00198 | |

### Supplementary Figure 12

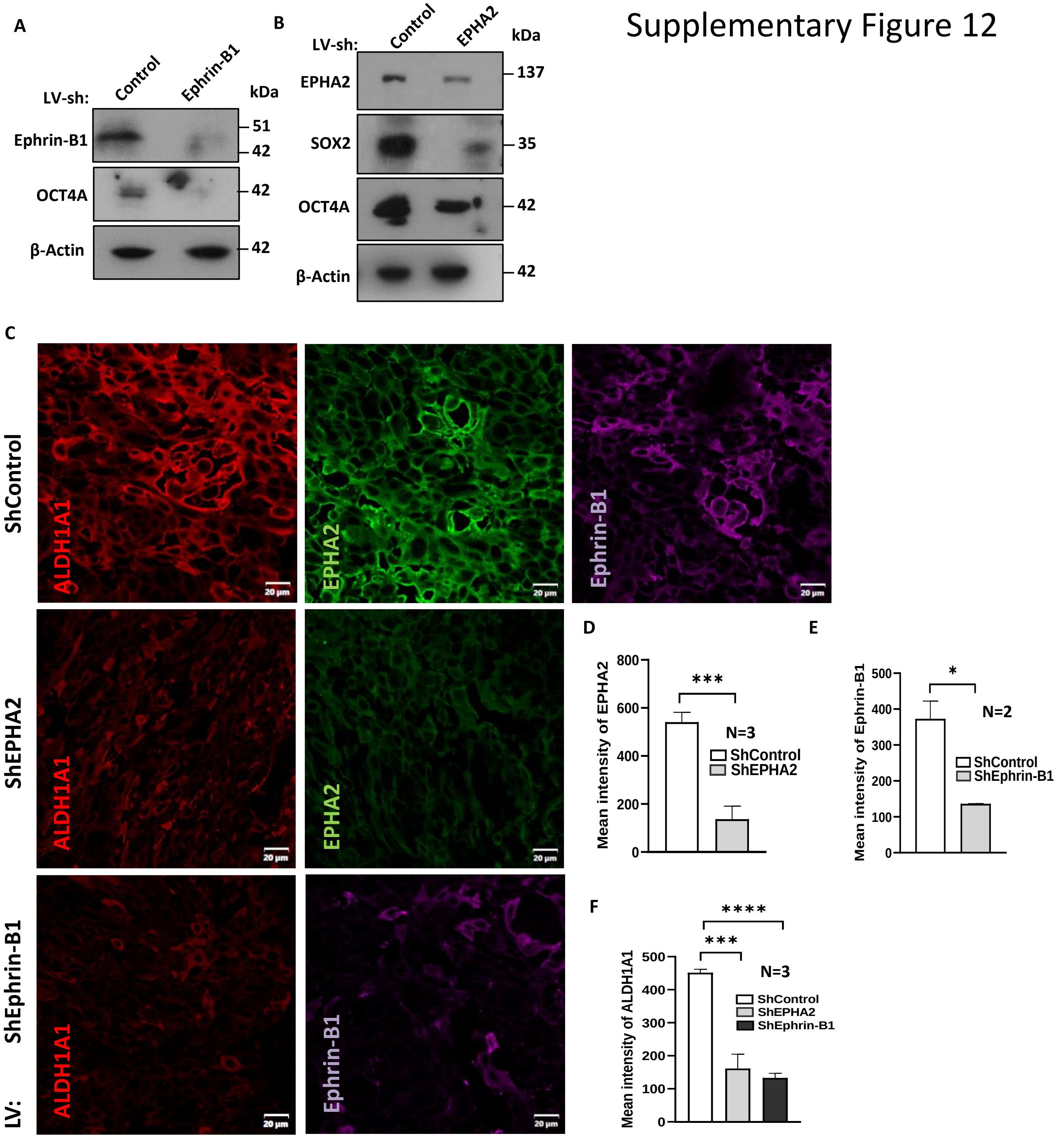

### Supplementary Figure 13

OSCC GSE13601

A

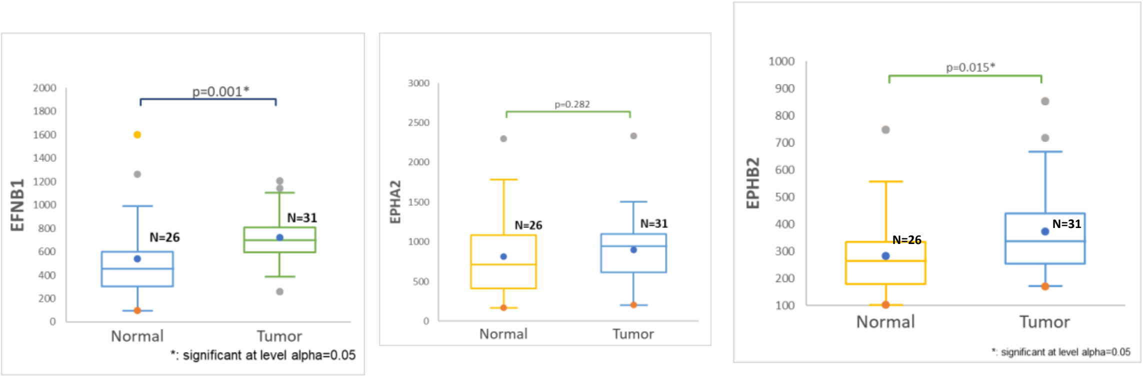


B

HNSCC TCGA; n=515

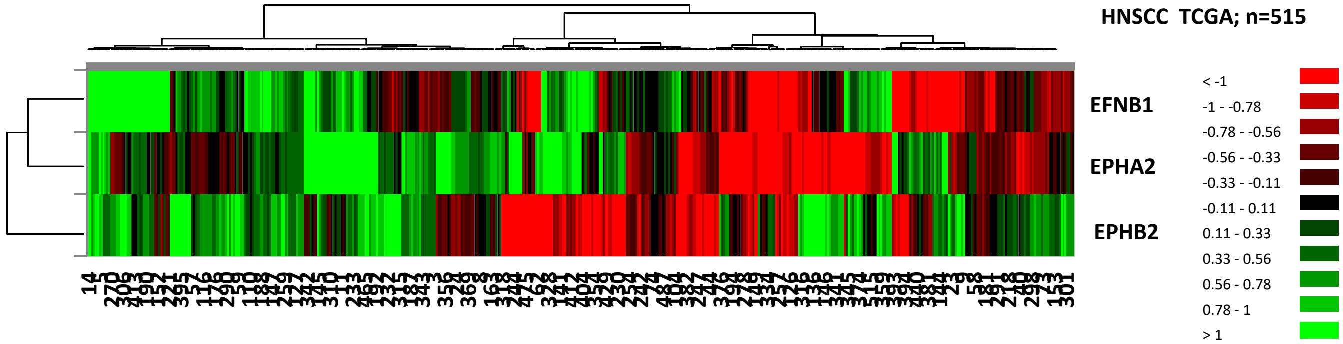


C

| HNSCC TCGA; n=515 | | |
|---|---|---|
| Correlation Matrix (Pearson): | | |
| Variables | EFNB1 | EPHA2 |
| EFNB1 | 1 | 0.283 |
| P value < 0.0001 | | |

| HNSCC GSE65858; n=255 | | |
|---|---|---|
| Correlation Matrix (Pearson): | | |
| Variables | EFNB1 | EPHA2 |
| EFNB1 | 1 | 0.235 |
| P value = 0 | | |
